# Defining the lipidome of Arabidopsis leaf mitochondria: Specific lipid complement and lipid biosynthesis capacity

**DOI:** 10.1101/2022.07.14.500104

**Authors:** Yi-Tse Liu, Jennifer Senkler, Cornelia Herrfurth, Hans-Peter Braun, Ivo Feussner

**Affiliations:** University of Goettingen, Albrecht-von-Haller-Institute for Plant Sciences, Department of Plant Biochemistry, 37077 Goettingen, Germany; Leibniz Universität Hannover, Institute of Plant Genetics, Department of Plant Proteomics, 30419 Hannover, Germany; University of Goettingen, Goettingen Center for Molecular Biosciences (GZMB), Service Unit for Metabolomics and Lipidomics, 37077 Goettingen, Germany; University of Goettingen, Goettingen Center for Molecular Biosciences (GZMB), Department of Plant Biochemistry, 37077 Goettingen, Germany

**Keywords:** *Arabidopsis thaliana*, lipidome, mitochondrium, proteome, respiration

## Abstract

Mitochondria are often considered the power stations of the cell, playing critical roles in various biological processes such as cellular respiration, photosynthesis, stress responses and programmed cell death. To maintain the structural and functional integrities of mitochondria, it is crucial to achieve a defined membrane lipid composition between different lipid classes wherein specific proportions of individual lipid species are present. Although mitochondria are capable of self-synthesizing a few lipid classes, many phospholipids are synthesized in the endoplasmic reticulum and transferred to mitochondria via membrane contact sites, as mitochondria are excluded from the vesicular transportation pathway. However, knowledge on the capability of lipid biosynthesis in mitochondria and the precise mechanism of maintaining the homeostasis of mitochondrial lipids is still scarce. Here we describe the lipidome of mitochondria isolated from *Arabidopsis* leaves, including the molecular species of glycerolipids, sphingolipids and sterols to depict the lipid landscape of mitochondrial membranes. In addition, we define proteins involved in lipid metabolism by proteomic analysis and compare our data with mitochondria from cell cultures since they still serve as model system. Proteins putatively localized to the membrane contact sites are proposed based on the proteomic results and online databases. Collectively, our results suggest that leaf mitochondria are capable - with the assistance of membrane contact site-localised proteins - of generating several lipid classes including phosphatidylethanolamines, cardiolipins, diacylgalactosylglycerols and free sterols. We anticipate our work to be a foundation to further investigate the functional roles of lipids and their involvement in biochemical reactions in plant mitochondria.

**One sentence summary:** The lipid landscape of plant mitochondria suggests that they are capable in generating several phospholipid classes with the assistant of membrane contact site-localized proteins.

## Introduction

Mitochondria are considered as semiautonomous organelles. According to a widely accepted hypothesis, they descend from proteobacteria that have been engulfed to form an eukaryotic cell. The two involved cells established an endosymbiosis (Gray et al., 1999). Mitochondria play crucial roles in various cellular processes, including ATP generation, metabolite biosynthesis, stress responses like phosphate starvation and initiation of programmed cell death (Jacoby et al., 2012; Michaud et al., 2016). The mitochondria of plant cells even have extended functions, many of which are related to photosynthesis (Braun, 2020). Mitochondria are enclosed by two membranes, the outer (OM) and the inner (IM) mitochondrial membranes, that both require defined protein and lipid compositions to maintain their functional integrity (Horvath and Daum, 2013; Michaud et al., 2017). The majority of the mitochondrial proteins is encoded by the nuclear genome, synthesized in the cytoplasm and post-translationally imported into the organelle. At the same time, a few proteins are encoded by the mitochondrial genome and synthesized within the organelle. Recent studies have broadened the knowledge of protein and lipid import in mitochondria, to the OM, the intermembrane space, the IM and the matrix (Michaud et al., 2017).

Biological membranes are composed of a wide variety of lipid classes and molecular species. According to the backbones of the lipid molecules, they are classified into three main lipid categories – glycerolipids, sphingolipids and sterols. To delineate the molecular species in this study, we later use the taxonomy containing two colon-separated units for the numbers of carbons and double bonds in the fatty acyl moiety, and an additional unit after a semicolon for the number of hydroxyl groups (when present) (Liebisch et al., 2020). Plant mitochondrial membranes contain high amounts of glycerophospholipids, such as phosphatidylcholine (PC) and phosphatidylethanolamine (PE), but contain less than 2 % of sterols (Moreau et al., 1974; Bligny and Douce, 1980). Sphingolipids have so far not been detected in mitochondrial membranes form plants. Most of these lipids are produced in the ER and are sorted to mitochondria and other organelles afterwards. Nevertheless, mitochondria have also their own capacity to generate some specific lipid classes. For instance, cardiolipin (CL) is synthesized in the IM and is present exclusively in mitochondria (Douce et al., 1972; Babiychuk et al., 2003). CL plays an essential role establishing the cristae of the IM and in maintaining the mitochondrial ultrastructure. CL is formed through the condensation of phosphatidylglycerol (PG) and diacylglycerol (DAG) and finally consists of four acyl chains. The inner envelope of the plastid is the major site for generating PG molecules (Müller and Frentzen, 2001). However, enzymes involved in PG biosynthesis are identified in mitochondria as well, suggesting that mitochondria are capable of self-synthesizing PG and thus CL (Babiychuk et al., 2003). In plants, the PG synthesizing enzymes, phosphatidylglycerolphosphate (PGP) synthase and PGP phosphatase (PGPP), are associated with mitochondria, and plastids; whereas CL synthase (CLS) localizes exclusively in the IM (Xu et al., 2002; Katayama et al., 2004). Notably, although mitochondrial lipid biosynthesis is not the major lipid source in plant cells, it is critical for certain organisms. In yeast, mitochondria are the major supplier of PE (Horvath and Daum, 2013). They generate PE from phosphatidylserine (PS) by phosphatidylserine decarboxylase (PSD). In *Arabidopsis,* three PSD enzymes have been identified and PSD1 localizes in mitochondria, providing PE molecules *in situ* (Nerlich et al., 2007). Alternatively, some PS molecules are converted from PE through the base-exchange pathway that substitutes the head groups of PE with serine molecules by PS synthase1, PSS1 (Yamaoka et al., 2011). While knowing mitochondria are capable of synthesizing PG, CL, PS and PE, its competence toward other lipid classes is still poorly understood (Li-Beisson et al., 2013).

Lipid trafficking between ER, mitochondria and plastids is essential for mitochondrial membrane biogenesis (Horvath and Daum, 2013; Michaud et al., 2017). Based on studies in yeast and mammals, glycerophospholipid biosynthesis takes place at a distinct membrane stretch of the ER, the mitochondria-associated membrane (MAM), wherein both ER and mitochondrial proteins have been identified (Vance, 1990). Elevated activities of PE, PC and PS synthesizing enzymes have been detected in purified yeast MAM. Moreover, this distinct membrane domain seems to occur ubiquitously among plants and animal cells, suggesting its critical role during evolution and in mediating lipid transfer between ER and mitochondria (Morré et al., 1971; Staehelin, 1997; Achleitner et al., 1999; Michaud et al., 2016). Although mitochondria and plastids work closely together in numerous pathways in plants, the lipid transport mechanism between these two organelles is largely unknown. Nevertheless, a few studies have suggested the relevance of lipid trafficking between these two organelles for survival, especially under environmental stresses. During phosphate starvation, higher numbers of mitochondria – plastid junctions are established (Jouhet et al., 2004). At the same time, drastic lipid remodeling of mitochondria, plastids and plasma membrane arises. PC and PE are degraded to release the phosphate residues for essential biological processes and the remaining molecules are recycled to generate the typical plastidial glyceroglycolipid, digalactosyldiacylglycerol (DGDG). This coincides with increased levels of CL in mitochondria isolated from *Arabidopsis* suspension cells and calli (Michaud et al., 2017). In *Arabidopsis,* the protein complex involved in lipid trafficking and tethering of the two mitochondrial membranes, the mitochondrial transmembrane lipoprotein (MTL) complex, has been identified recently (Michaud et al., 2016). MTL is composed of more than 200 subunits and it has been demonstrated that MTL promotes the translocation of PE from IM to OM and the import of DGDG from plastids to mitochondria during phosphate starvation. Furthermore, mutation of a newly identified MTL subunit, digalactosyldiacylglycerol synthase suppressor 1 (DGS1), leads to alteration of plastidial and mitochondrial lipid composition and deficiency in mitochondrial biogenesis (Li et al., 2019). The characterization of the MTL complex provides an initial insight in understanding the mechanism of lipid trafficking between mitochondria and plastids.

In this study, we aimed to characterize the lipid metabolism of *Arabidopsis* leaf mitochondria in depth and to compare these data with published and own data from suspension cell cultures since they serve as model system. Therefore, we conducted an in-depth lipidomic analysis, providing the molecular species information of all lipid categories including glycerolipids, sphingolipids and sterols to illustrate the lipid landscape of mitochondria in *Arabidopsis* leaves. In combination with a proteomic approach, we intended to specify the capacity of lipid biosynthesis and modification in plant mitochondria, defining lipid species and classes that may be generated by mitochondria. We additionally propose putative membrane contact site-localizing proteins and their roles in interorganelle communication. Our results suggest that leaf mitochondria possess a defined lipid composition wherein specific lipid molecular species appear. In addition, lipid trafficking between mitochondria, ER and plastids via membrane contact sites provides assistance in maintaining the homeostasis of the lipid composition in mitochondria.

## Results

### Purity of mitochondrial fractions

A combinatorial approach by lipidomics, proteomics and mining of online databases was applied to investigate the lipid composition as well as the capacity of lipid metabolism of *Arabidopsis* leaf mitochondria. Mitochondria were isolated from three independent experiments by differential centrifugation combined with Percoll density gradient centrifugation. To ensure and evaluate the purity of the mitochondrial fractions, all samples were investigated by two-dimensional (2D) blue native (BN)/SDS PAGE and by shotgun proteome analyses. Our mitochondrial fractions proved to be of high purity. The two photosystems were not detectable on our 2D gels. Only trace amounts of the large subunit from Rubisco, the most abundant protein in leaves, were visible in the leaf mitochondrial fractions (L-mito) on BN/SDS gels, and the small subunit of Rubisco was barely visible (Fig. 1A, Supplemental Fig. S1). Since most of the work on plant mitochondria was performed with suspension cell cultures so far, we isolated mitochondria from dark-grown *Arabidopsis* cell culture (C-mito) for comparison. They were investigated by the same procedure and used as an established control for contamination from chloroplasts. The L-mito and C-mito fractions were highly similar on BN/SDS PAGE (Fig. 1, Supplemental Fig. S1). In addition, the purity of the mitochondrial fractions was investigated by label-free quantitative shotgun proteomics. Building on the proteome data (Supplemental Tab. S1), the proportion of mitochondria-localized proteins was calculated to be in the range of 87 to 94 % (Fig. 1B, Supplemental Fig. S2). These results indicate that the mitochondrial fractions are of high purity.

**Figure 1.**
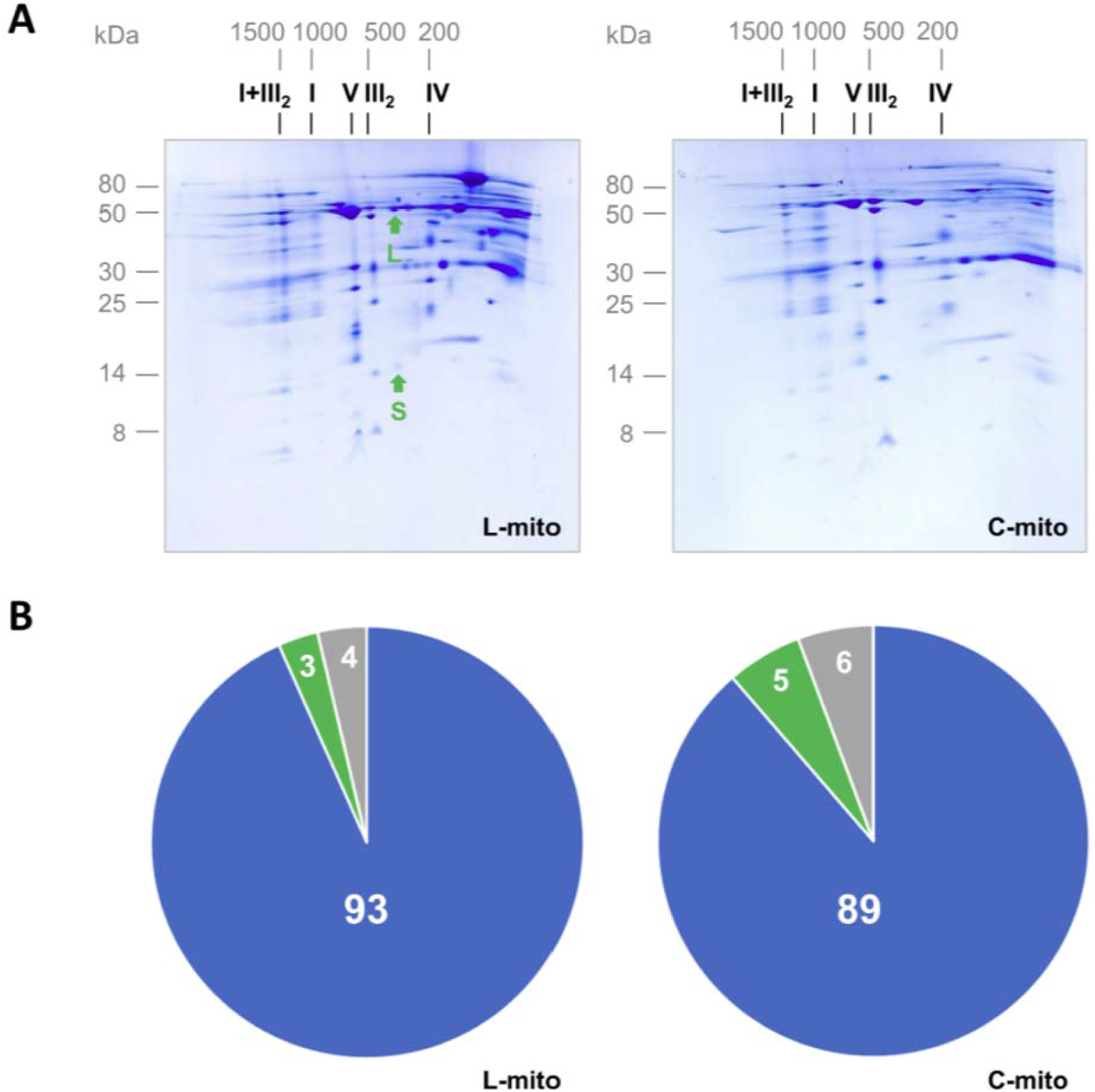
Purity of mitochondrial fractions. The purity of mitochondrial fractions was determined by 2D blue native / SDS PAGE **(A)** and by summed-up peptide intensities of subcellular compartments based on protein assignments as given by the Subcellular localization database for Arabidopsis proteins (SUBAcon; www.suba.live) **(B)**. **A**: Mitochondria were isolated from *Arabidopsis* leaves (L-mito) and cell cultures (C-mito). Proteins were separated by 2D Blue native PAGE and Coommassie-stained. Numbers on top and to the left of the 2D gels refer to the masses of standard protein complexes / proteins (in kDa), the roman numbers above the gels to the identity of OXPHOS complexes. I+III_2_: supercomplex consisting of complex I and dimeric complex III; I: complex I; V: complex V; III_2_: dimeric complex III; IV: complex IV. The small (S; 14.5 kDa) and the large (L; 53.5 kDa) subunit of Rubisco are indicated by green arrows. Mitochondrial preparations derive from three independent experiments, and the corresponding 2D gels of the two remaining biological replicates as well as one reference gel each for mitochondrial and chloroplast fractions from *Arabidopsis* cell culture (Mito ref) or leaves (Cp rev) are shown in Supplemental Fig. S1. These three preparations were used for lipidomics with LC-MS/MS (Fig. 3, 5, 6, and S3) and proteomics **(B)**. **B:** Mitochondrial fractions from *Arabidopsis* leaves and cell cultures were analyzed by label-free quantitative shotgun proteomics. Peptide intensities assigned to subcellular compartments were summed-up and averaged results for L-mitos and C-mitos from the three independent experiments were visualized by pie charts (for detailed results see Supplemental Fig. S2 and Supplemental Tab. S1). Blue: mitochondria; green: plastids; gray: others; numbers in %.

### PC, PE and CL are strongly enriched glycerolipids in plant leaf mitochondria

Glycerolipids are the most abundant lipids not only in mitochondria but also generally in plant tissues, accounting for more than 90 % of the overall lipids in leaves (Li-Beisson et al., 2013; Michaud et al., 2017). Glycerophospholipids are fundamental to all biological membranes, while glyceroglycolipids are critical for photosynthetic membranes and localize mainly in plastids of vegetative tissues (Hölzl and Dörmann, 2019). Therefore, lipids were extracted from leaves and purified mitochondria therefrom to determine the proportion of the major glycerolipid classes in a quantitative approach based on thin layer chromatography followed by gas chromatography coupled to flame ionization detection (TLC-GC/FID, Fig. 2). In L-TE, glyceroglycolipids including monogalactosyldiacylglycerols (MGDG; 38.9 ± 1.1 %) and DGDG (22.3 ± 3.8 %) contribute to the majority of the overall lipids; while glycerophospholipids, PC (35.7 ± 6.0 %) and PE (37.2 ± 0.3 %), are the most abundant lipids in L-mito. CL accounts for 4.0 ± 0.9 % in the lipidome of L-mito; however, it was not detectable in L-TE because of its low abundance in total cellular lipids. PG, as the precursor of CL, consists 5.6 ± 2.1 % of the overall lipids in L-mito.

**Figure 2.**
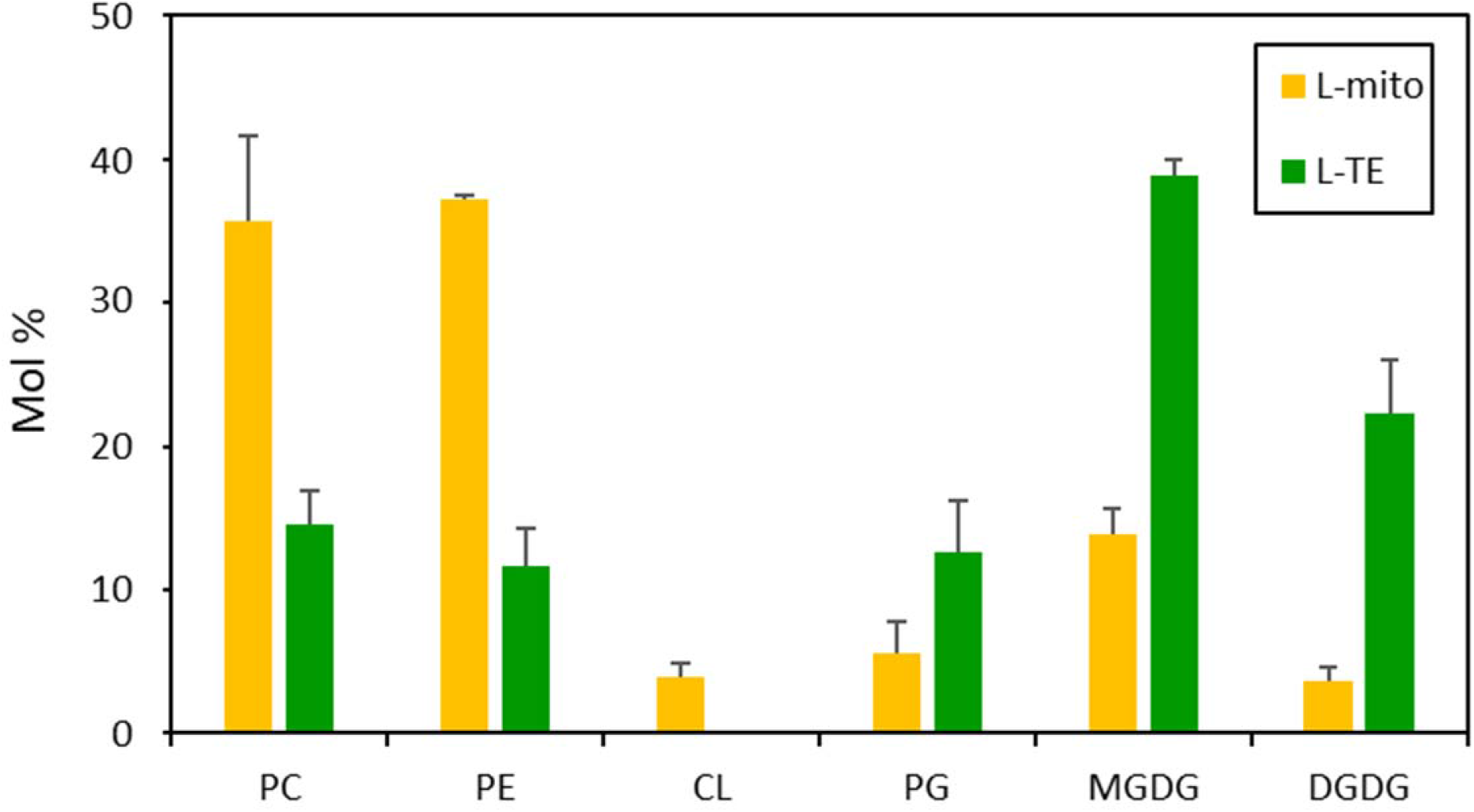
Lipid class profiles of purified mitochondria and total leaf extracts. Glycerolipids of leaf total extract (L-TE) and mitochondria isolated from leaves (L-mito) were analyzed quantitatively by a TLC-GC/FID approach. Data of L-TE represent mean values in mol % from three independent experiments ± SD; data of L-mito represent mean values in mol % from two of the three independent experiments ± SEM. PC, phosphatidylcholine; PE, phosphatidylethanolamine; CL, cardiolipin; PG, phosphatidylglycerol; MGDG, monogalactosyldiacylglycerol; DGDG, digalactosyldiacylglycerol.

### Specific molecular glycerolipid species are enriched in plant leaf mitochondria

Glycerophospholipids are the most abundant lipids in mitochondria (Fig. 2) (Horvath and Daum, 2013; Michaud et al., 2017). Therefore, the molecular species of glycerophospholipids, as well as those of glyceroglycolipids and diacylglycerols, were analyzed from the same samples originating from three independent experiments as for proteomics (Fig. 1) by LC-MS/MS (Fig. 3a). Data are expressed in mol %. The most abundant PC and PE molecular species in mitochondria with the acyl chains 18:2/18:3, 18:2/18:2 and 18:3/18:3 account for more than 60 % in both lipid classes (Fig. 3a; Supplemental Tab. S2). The species 16:0/18:2, 16:0/18:3 of both lipid classes however were depleted by about 50 % in this organelle in comparison to L-TE. PS (18:0/18:2) composes up to 38.3 % in L-mito with an increase of 2^4.3^ fold comparing to L-TE. The accumulation of this lipid species was the most remarkable one that was even visible when the fatty acid profiles of the different lipid classes were calculated from the molecular species data, showing an enrichment of 18:0 in mitochondria on the expense of 16:0 (Supplemental Fig. S3). Together the main molecular species of these three lipid classes have a C18/C18 acyl chain composition in common.

**Figure 3.**
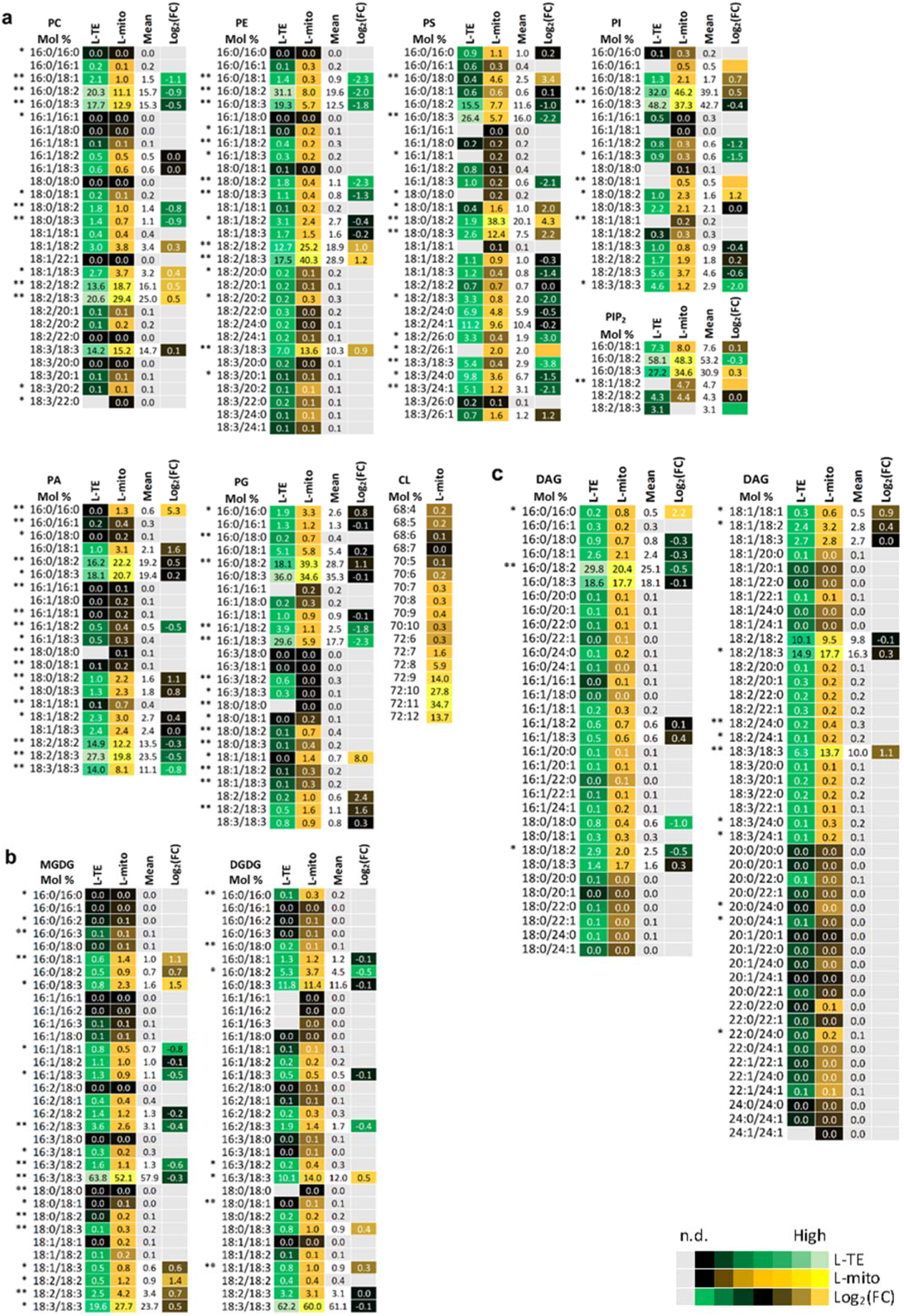
Profiles of the distribution and fold changes of the molecular glycerolipid species between L-mito and L-TE. Heat map visualizations of (a) glycerophospholipids, (b) glyceroglycolipids and (c) diacylglycerols illustrate the difference of species distribution based on LC-MS/MS analyses. Each lipid class is represented by one set of joined columns, only DAG is divided into two sets due to space constraints. Identity of columns in each set from left to right: Column 1 lists the individual lipid class and the identity of the detected molecular lipid species. Column 2 (L-TE) lists the respective distribution of each molecular species in L-TE, expressed as the mean of its relative values in mol % in the three independent experiments also used for proteomics and for sphingolipid and sterol analysis (Figs. 1, 5 and 6, Supplemental Tab. S1). Column 3 (L-mito) lists the respective distribution of each molecular species in L-mito, expressed as the mean of its relative values (mol %) in the three independent experiments also used for proteomics. Column 4 lists the mean distribution of both sample types, and column 5 (Log_2_(FC)) lists the binary logarithm of fold change. Binary logarithm was applied when the mean values are higher than 0.5 to inspect the fold changes (FC) between L-mito and L-TE. The heat map colors represent mean intensity values according to the color map on the low right-hand side. One and two asterisks (*, **) indicate p values < 0.05 and < 0.01, respectively, by Student’s t-test. PC, phosphatidylcholine; PE, phosphatidylethanolamine; PS: phosphatidylserine; PI: phosphatidylinositol; PA phosphatidic acid; DAG: Diacylglycerol; CL, cardiolipin; PG, phosphatidylglycerol; MGDG, monogalactosyldiacylglycerol; DGDG, digalactosyldiacylglycerol.

An important intermediate in glycerolipid metabolism is phosphatidic acid (PA), serving as precursor or product for numerous glycerolipids including PC, PE, PG, phosphatidylinositol (PI) and DAG. PA (16:0/18:2), PA (16:0/18:3) and PA (18:2/18:3) are the most abundant PA species in both fractions, L-mito and L-TE. Notably, only a minor molecular PA species (16:0/16:0; 1.3 mol %) accumulates preferentially in mitochondria with an enrichment of 2^5.3^ fold comparing to L-TE. DAG as the other important lipid intermediate displays a higher complexity in its species profile. DAG (16:0/18:2), DAG (16:0/18:3), DAG (18:2/18:3) and DAG (18:3/18:3) build a substantial amount (>65 mol %) of the DAG profile in L-mito (Fig. 3c). As in case of PA, only the minor DAG species (16:0/16:0) shows a 2^2.2^ fold increase in L-mito compared to L-TE.

PG is formed from PA in the envelopes of plastids and mitochondria and serves as precursor for the mitochondrial lipid CL (Babiychuk et al., 2003). Hydrolysis and condensation of PG lead to the formation of DAG and CL correspondingly. PG (16:0/18:2) and PG (16:0/18:3) are the major species in L-mito, but only the former species is enriched in mitochondria (2^1,1^ fold). In addition, unsaturated PG species are significantly enriched in L-mito, which contribute to the structures of the down-stream CL molecules. CL (72:10) and CL (72:11), composed of 18:2 and 18:3 acyl chains, are the most abundant CL (Fig. 3a).

Glycerophospholipids serve not only as membrane building blocks but are also important precursors for signaling molecules in the cell. PI, phosphatidylinositol monophosphate (PIP) and phosphatidylinositol bisphosphate (PIP_2_) exert regulatory functions in cell development and polarity determination (Heilmann, 2016). In L-mito, molecular species with 16:0/18:2 and 16:0/18:3 acyl chains compose up to 80 % of PI and PIP_2_; yet PIP was not detectable in L-mito. PI (16:0/16:1) and PI (18:0/18:1) were only detectable in the mitochondrial extracts, and PI (18:0/18:2) is significantly enriched in L-mito for 2^1.2^ fold in comparison to L-TE (Fig. 3a). On the other hand, we obtained higher signals of PIP_2_ (18:1/18:2) in L-mito.

Glyceroglycolipids carry carbohydrate residues as their head groups; for instance, the galactose-containing lipids, MGDG and DGDG contain one and two galactoses, respectively (Hölzl and Dörmann, 2019). MGDG (16:3/18:3) and DGDG (18:3/18:3) are the major components of the overall glyceroglycolipids in both L-TE and L-mito (Fig. 3b). The amount of MGDG (16:0/18:3) and MGDG (18:2/18:2) are specifically elevated in L-mito for 2^1.5^ and 2^1.4^ folds, respectively, compared to L-TE. In contrast, the species MGDG (16:1/18:1) and MGDG (16:3/18:2) are significantly decreased in L-mito. In summary, the major mitochondrial lipids PC, PE, PS and CL consist primarily of C18/C18 species, while PI, PIP_2_ and PG consist primarily of C16/C18 species and the lipid precursors DAG and PA resemble a mixture of both backbones.

### Proteomic analyses provides insights into the leaf mitochondrial glycerolipid metabolism capacity

To further analyze the proteome of the mitochondrial fractions listed in Supplemental Tab. S1, their protein composition was investigated for enzymes from lipid metabolism. Subsequently, their biological function and the subcellular localization were retrieved from The Arabidopsis Information Resource (TAIR) and the Subcellular localization database for Arabidopsis proteins (SUBAcon) databases (Supplemental Fig. S4).

About 40 proteins involved in the biosynthesis and modification of fatty acids and more complex lipids were identified in mitochondrial extracts (Supplemental Tab. S3). Four enzymes from glycerophospholipid metabolism were detected in all mitochondrial fractions (Fig. 4). Two glycerol-3-phosphate dehydrogenases were identified (SDP6 and GPDHp), being involved in an early step of glycerophospholipid biosynthesis. Phosphoethanolamine cytidylyltransferase (PECT) was detected in our study by similar abundances in the L-mito and C-mito samples. CL is synthesized after the condensation of PG and CDP-DAG by CL synthase (CLS) or after the transacylation of monolyso-CL by monolysocardiolipin acyltransferase (LCLAT or Tafazzin)(Xu et al., 2006). CLS and Tafazzin were identified in the shotgun proteomic analysis in this study with a higher abundance in C-mito compared to L-mito. PG is synthesized from PA via CDP-DAG by phosphatidylglycerophosphate synthase 1 (PGP1) (Xu et al., 2002) and it was only detected in C-mito samples. Phosphatidylserine decarboxylase 1 (PSD1) synthesizes PE by decarboxylating the headgroup of PS (Birner et al., 2001; Nerlich et al., 2007). Again in our study PSD1 was only detected in C-mito samples. Interestingly PE and PC can be interconverted into each other via PA and the detected phospholipase Dα1 (PLDα1) enzyme activity. PLDα1 was preferentially found in C-mito samples. PIP and PIP_2_ can be degraded by *myo*-inositol polyphosphate 1-phosphatase (SAL1). Although many enzymes involved in phosphoinositide metabolism have been annotated to localize in mitochondria, only SAL1 was identified in our approach in the C-mito samples.

**Figure 4.**
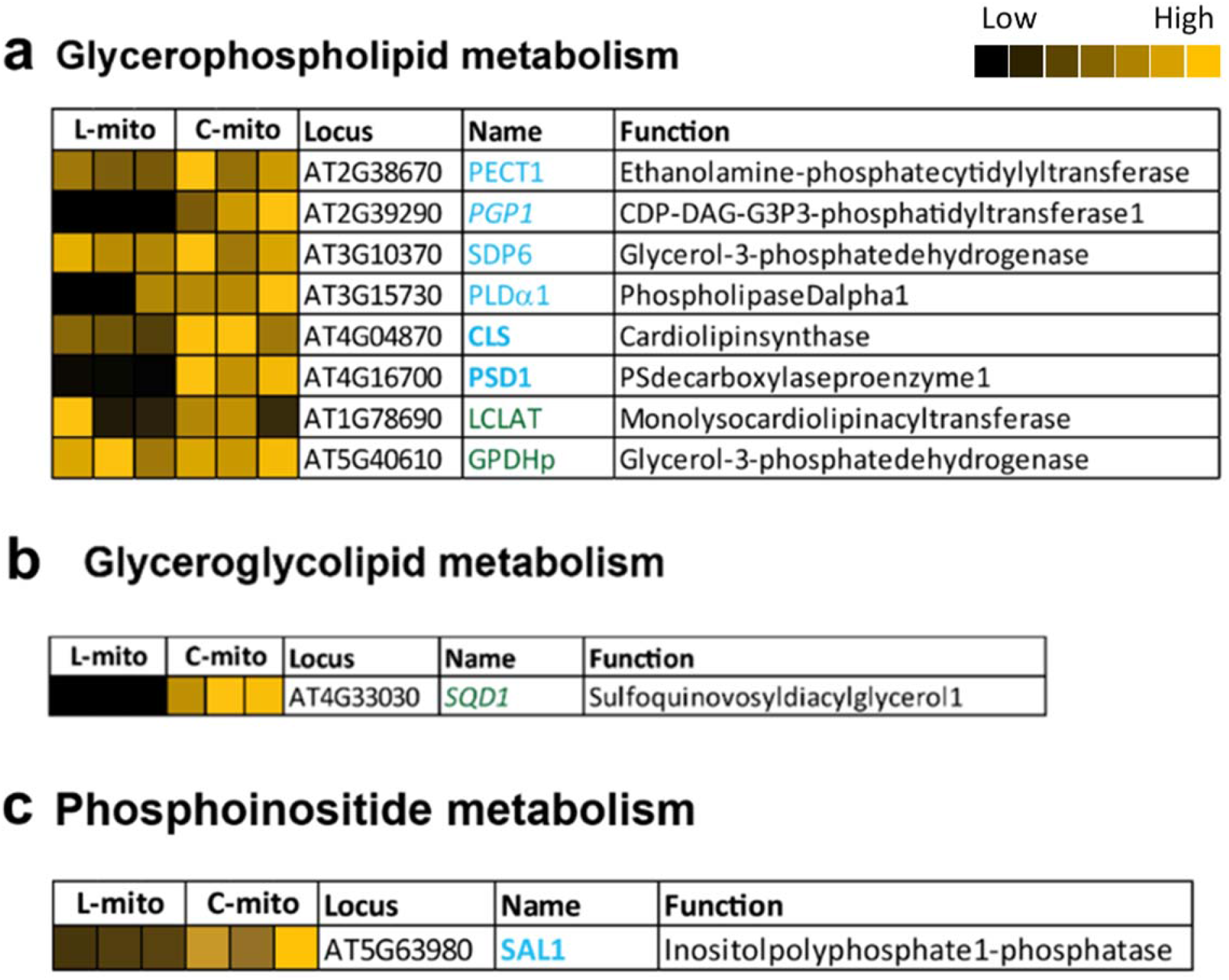
Mitochondrial localized proteins that are related to glycerolipid biosynthesis. (a) Glycerophospholipid, (b) glyceroglycolipid and (c) phosphoinositide metabolism. Proteins identified and/or localized in mitochondria were labeled (i) blue: proteins identified in the proteomic analysis of this study and also predicted to localize in mitochondria, (ii) green: proteins identified in the proteomic analysis of this study in mitochondria but predicted to localize in other organelles, (iii) bold font: exclusively localized in mitochondria and (iv) italic font: only identified in one of the mitochondrial populations. Heat maps visualize the protein abundance of three independent experiments of mitochondria purified from leaves and cell cultures from Fig. 1 and listed in Supplemental Tabs. S1 and S3. The heat map colors represent mean intensity values according to the color map on the top right-hand side. Predicted protein localization was based on The Arabidopsis Information Resource (TAIR; www.arabidopsis.org) and the Subcellular localization database for Arabidopsis proteins (SUBAcon; www.suba.live).

The biosynthesis of glyceroglycolipids takes place in the plastids envelopes by MGDG and DGDG synthases (MGDs and DGDs), respectively (Hölzl and Dörmann, 2019). MGDG is generated by adding a galactose head group to DAG; subsequent addition of another galactose by DGD1 generates DGDG. The detected specific species profiles of mitochondrial glyceroglycolipids may suggest at least for a specific transport from plastids to mitochondria (Fig. 3b). The sulfolipid SQDG is synthesized in two steps. Uridine diphosphate (UDP)-glucose is first combined with a sulfite by UDP-sulfoquinovose synthase 1 (SQD1), followed by transferring the sulfoquinovose head group to DAG and thus forms SQDG. Although SQD1 was identified in the C-mito samples in this study, we did not detect SQDG lipids in any of the mitochondrial samples. Together we could exclusively localize three biosynthetic steps to mitochondria: (i) CLS and catalyzing the formation of CL as well as (ii) PSD1 and (iii) PECT1 facilitating PE formation.

### Specific molecular sphingolipid and sterol species suggest a function in mitochondria

Sphingolipids and sterols are important modulators of membrane microdomains and play critical roles in regulating the balance between cell survival and apoptosis. Therefore, we profiled next their mitochondrial compositions in comparison to the total lipidome of leaves in the same samples that were used for proteomics and glycerolipid analysis (Figs. 1 and 3) by LC-MS/MS (Figs. 5 and 6) and expressed the lipid profiles in mol %. For sphingolipids both simple and complex sphingolipids were measured, including long-chain bases (LCB), phosphorylated LCB (LCB-P), ceramides (Cer), glucosylceramides (GlcCer) and glycosyl inositol phosphoceramides (GIPC) (Fig. 5). All sphingolipids have LCBs, 18-carbon amino-alcohols, as their backbones. Phytosphingosine (18:0;3), hosting three hydroxyl groups, is the most abundant free LCB in both L-mito and L-TE (Fig. 5a). However, dihydrosphingosine (18:0;2) is highly enriched in L-mito as well as phosphorylated dihydrosphingosine (18:0;2-P), which is the major component in the LCB-P pool of L-mito (64.7 %). LCBs can be further *N*-acylated to generate Cer, the basic structure of complex sphingolipid classes. Addition of glucoses to Cer generates GlcCer, following sequential extension of phosphoinositol, hexose and/or hexose derivatives generate series of GIPCs. Series 0, A and B GIPCs carry one, two and three additional hexoses on the head groups of inositol phosphoceramides, respectively (Cacas et al., 2012; Haslam and Feussner, 2022). Similar to LCB and LCB-P, only specific species were enriched in L-mito (Fig. 5b): Cer (18:0;3/24:0;1), GlcCer (18:1;2/24:1;1) and series A hexose-carrying GIPC (H-GIPC) (18:1;3/24:1;1). Remarkably, GIPCs were only detectable in mitochondrial samples (both L-mito and C-mito) in our approach. However, proteins related to sphingolipid metabolism were neither identified in plant mitochondria via proteomic analyses, nor retrieved from online databases.

**Figure 5.**
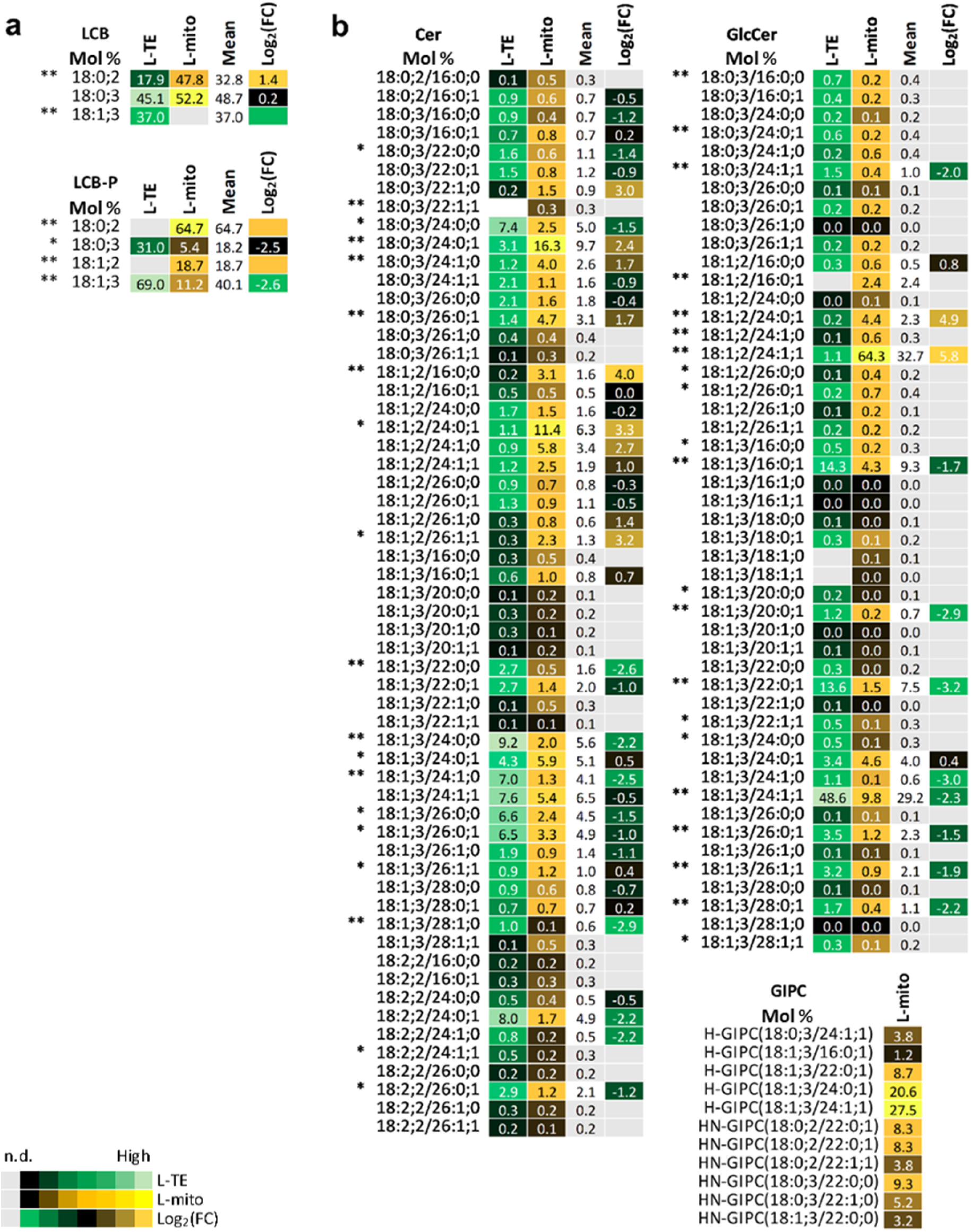
Profiles of the distribution and fold changes of the molecular sphingolipid species between L-mito and L-TE. Heat map visualizations of (a) long chain bases, long chain base-phosphates and (b) complex sphingolipids illustrate the difference of species distribution based on LC-MS/MS analyses. Each lipid class is represented by one set of joined columns. Identity of columns in each set from left to right: Column 1 lists the individual lipid class and the identity of the detected molecular lipid species. Column 2 (L-TE) lists the respective distribution of each molecular species in L-TE, expressed as the mean of its relative values (mol %) in the three independent experiments also used for proteomics and for glycerolipid and sterol analysis (Figs. 1, 3 and 6, Supplemental Tab. S1). Column 3 (L-mito) lists the respective distribution of each molecular species in L-mito, expressed as the mean of its relative values (mol %) in the three independent experiments also used for proteomics. Column 4 lists the mean distribution of both sample types, and column 5 (Log_2_(FC)) lists the binary logarithm of fold change. Binary logarithm was applied when the mean values are higher than 0.5 to inspect the fold changes (FC) between L-mito and L-TE. The heat map colors represent mean intensity values according to the color map on the low left-hand side. One and two asterisks (*, **) indicate p values < 0.05 and < 0.01, respectively, by Student’s t-test. LCB, long-chain base; LCB-P, long chain base-phosphate; Cer, ceramide; GlcCer, glycosylceramide; GIPC, glycosyl inositol phosphoceramide.

**Figure 6.**
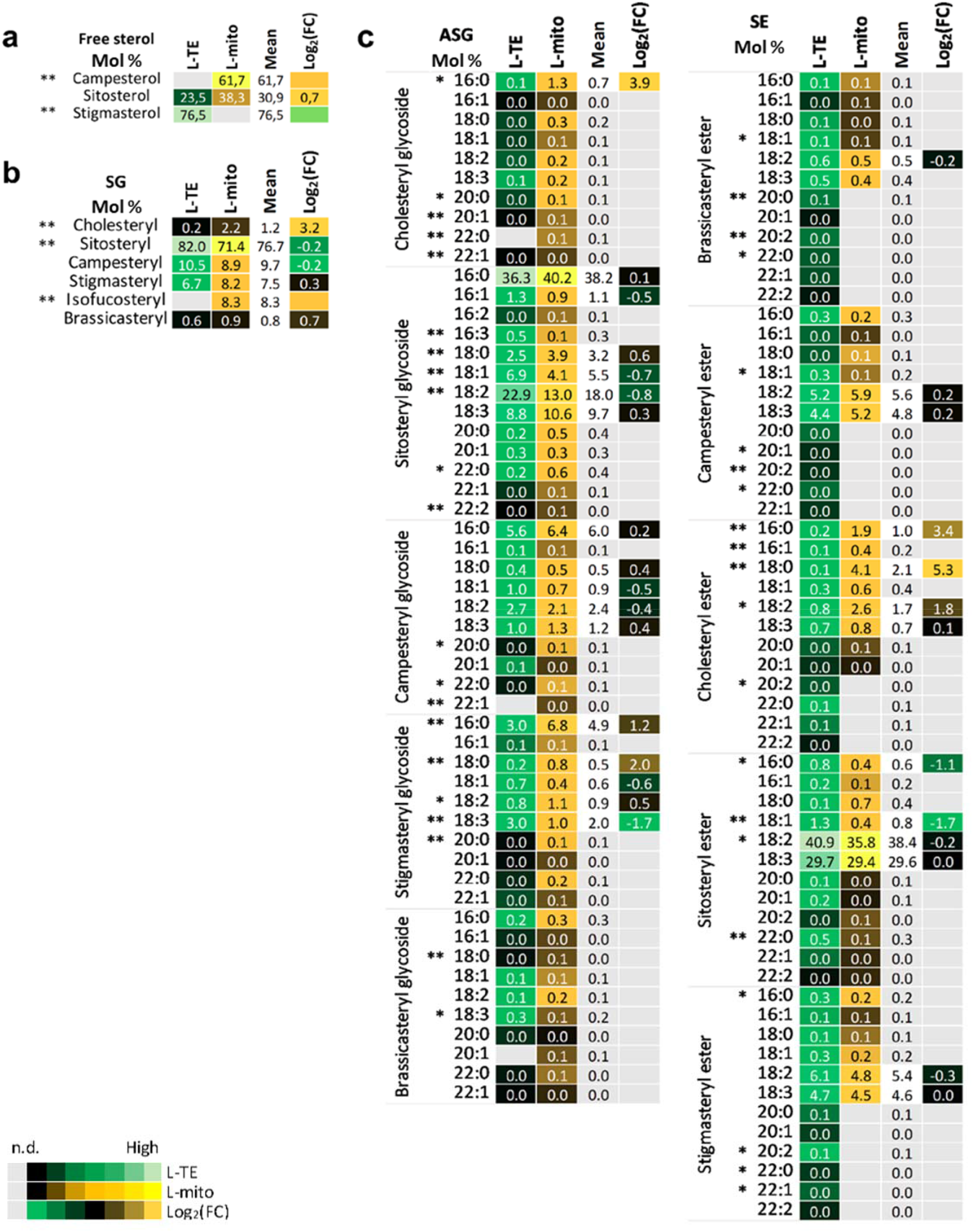
Profiles of the distribution and fold changes of the molecular sterol species between L-mito and L-TE. Heat map visualizations of (a) free sterols (b) steryl glycosides (SG) and (c) acylated steryl glycosides (ASG) and steryl esters (SE) illustrate the difference of species distribution based on LC-MS/MS analyses. Each lipid class is represented by one set of joined columns. Identity of columns in each set from left to right: Column 1 lists the individual lipid class and the identity of the detected molecular lipid species. Column 2 (L-TE) lists the respective distribution of each molecular species in L-TE, expressed as the mean of its relative values (mol %) in the three independent experiments also used for proteomics and for glycerolipid and sphingolipid analysis (Figs. 1, 3 and 5, Supplemental Tab. S1). Column 3 (L-mito) lists the respective distribution of each molecular species in L-mito, expressed as the mean of its relative values (mol %) in the three independent experiments also used for proteomics. Column 4 lists the mean distribution of both sample types, and column 5 (Log_2_(FC)) lists the binary logarithm of fold change. The heat map colors represent mean intensity values according to the color map on the low left-hand side. Binary logarithm was applied when the mean values are higher than 0.5 to inspect the fold changes (FC) between L-mito and L-TE. One and two asterisks (*, **) indicate p values < 0.05 and < 0.01, respectively, by Student’s t-test.

The common structure of sterols is a four-ring system, cyclopentanoperhydrophenanthrene, with possible conjugation of hydroxyl groups and acyl chains. In plants, a complex mixture of sterols can be found, including brassicasterol, campesterol, cholesterol, sitosterol and stigmasterol (Cacas et al., 2012). Campesterol was identified as major free sterol with 61.7 % in L-mito (Fig. 6). From the group of steryl glycosides cholesteryl and isofucosteryl glycoside accumulated preferentially in L-mito. Considering SE, 16:0 and 18:0 containing cholesteryl esters were enriched 2^3.4^ and 2^5.3^ folds, respectively, in L-mito. For ASG however the situation was blurred. DWF1 was the only protein related to sterol metabolism that we identified in plant mitochondria via proteomic analyses. In summary, we observed an accumulation of GIPCs and free campersterol in leaf mitochondria.

Mitochondria harbor various additional metabolic pathways for the synthesis of lipophilic molecules. More than 20 fatty acid biosynthesis-related proteins were identified in our mitochondrial proteome (Supplemental Fig. S5). They contribute to the synthesis of lipoic acid, ubiquinone and other terpenoid-quinones. Abundance, function and predicted localization of these proteins are specified in Supplemental Tab. S1 and S3.

### Lipid molecules can be imported by mitochondria

Transport of DGDG from plastids to mitochondria and reallocation of PE from IM to OM via the MTL complex have been demonstrated under phosphate-depleting conditions (Michaud et al., 2016). The MTL complex has more than 200 subunits and 11 have been verified to associate physically with the core subunits, Mic60 and Tom40, as previously shown by immunoblotting (Michaud et al., 2016; Li et al., 2019). In our mitochondrial proteome listed in Supplemental Tab. S1, 186 of the subunits (87 %) were identified including all the 11 verified ones (Fig. 7, Supplemental Tab. S4). The newly identified MTL subunit, DGS1, which links mitochondrial protein to lipid transport, is present in all mitochondrial extracts.

**Figure 7.**
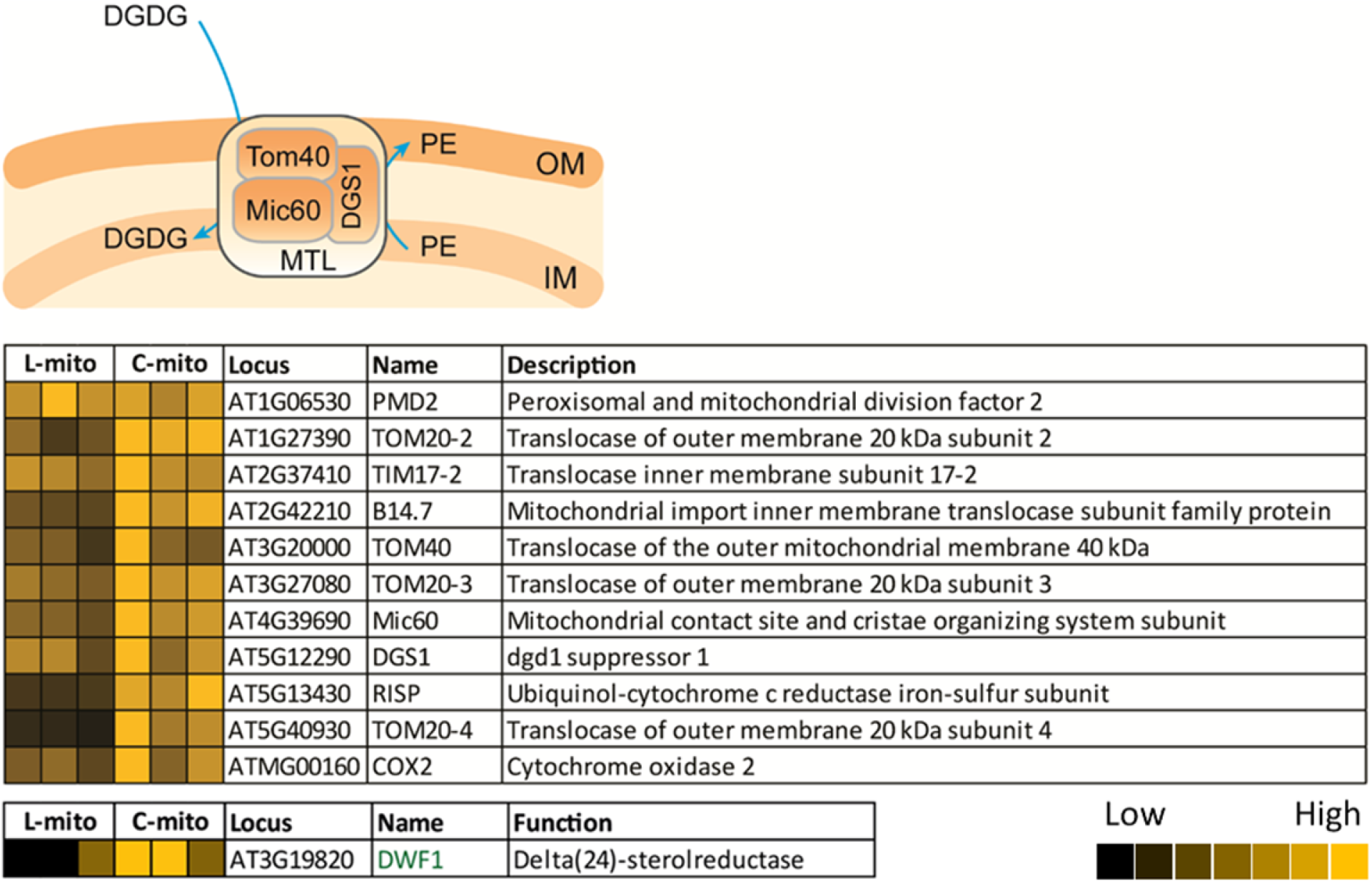
MTL complex-associated proteins in mitochondria. Our proteomic analyses covered 186 of the 214 hypothetical subunits in the MTL complex (listed in Supplemental Tab. S4); 11 have been investigated by immunoblotting approaches in previous studies. Heat maps visualize the protein abundance of three independent experiments of mitochondria purified from leaves and cell cultures. The heat map colors represent mean intensity values according to the color map on the low right-hand side.

In addition, at least 10 outer envelope (OE)-localized proteins from plastids were co-purified with mitochondria, with higher abundance in C-mito (Fig. 8, Supplemental Tab. S5). They account for 2.1 % (10/471) of the identified plastidial proteins, which are enriched in comparison to the proportion of OE-localized proteins in plastids (46/3002, 1.5 %) (Inoue, 2007, 2011; Simm et al., 2013; Kim et al., 2019). The enrichment of OE-localized proteins in mitochondrial extracts suggest a close physical interaction between mitochondria and plastidial OE membrane. For instance, the membrane contact sites, which were not disrupted during the isolation procedure, and therefore are present in the mitochondrial samples as membrane patches.

**Figure 8.**
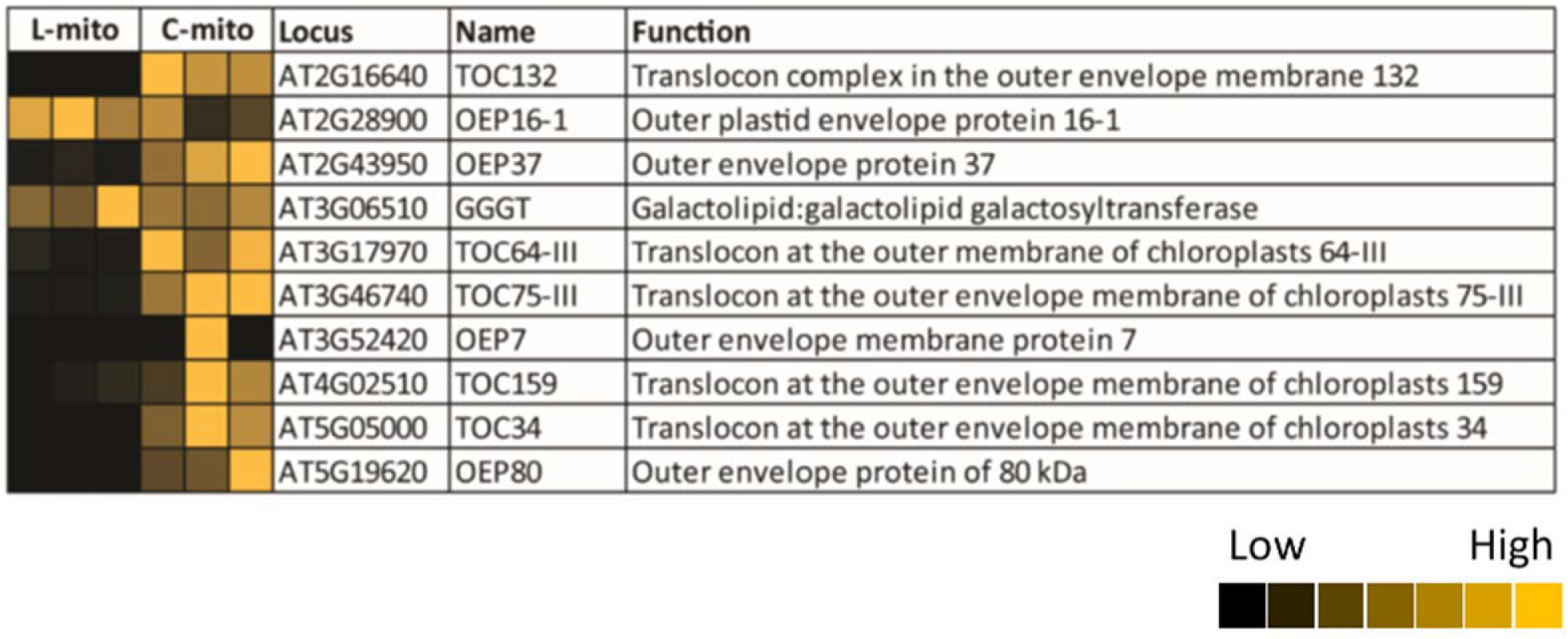
Identified plastid outer envelope proteins in mitochondrial extracts. Our proteomic analyses identified 10 outer envelope-localized proteins (listed in Supplemental Tab. S5). Heat maps visualize the protein abundance of three independent experiments of mitochondria purified from leaves and cell cultures. The heat map colors represent mean intensity values according to the color map on the left-hand side.

## Discussion

In this study, we present an in-depth dataset of lipid molecular species from glycerolipids, sphingolipids and sterols of Arabidopsis leaf mitochondria. With the assistance of online resources, we assigned and confirmed central lipid biosynthesis steps within these mitochondrial fractions. Furthermore, we confirmed and expanded our knowledge on the existence of a protein complex for lipid trafficking between mitochondria and other organelles. In the past, the most abundant plant mitochondrial glycerolipid classes, such as PC, PE, PI, PA, CL, MGDG and DGDG, from *Arabidopsis* cell cultures and calli under phosphate-depleted conditions have been quantified with TLC-GC (Jouhet et al., 2004; Michaud et al., 2016). However, a TLC-GC-based approach lacks the molecular species information concerning acyl chain length and unsaturation degree of the lipids, which are critical parameters to determine the membranes physical properties. In addition, neither sphingolipids nor sterols of mitochondria were profiled to our very best knowledge formerly in *Arabidopsis.*

In our study, mitochondria were purified from leaves by differential and Percoll density gradient centrifugations and compared with the well established mitochondrial preparations from suspension cell culture (Jouhet et al., 2004; Klodmann et al., 2011; Behrens et al., 2013; Michaud et al., 2016). Much effort was made to document purity of the resulting organelle fraction: (i) visual inspection of the protein complex composition by 2D BN / SDS PAGE and (ii) label-free quantitative shotgun proteins in combination with SUBAcon evaluation. The SUBAcon database integrates worldwide knowledge on subcellular localization information of Arabidopsis proteins based on *in vitro* or *in vivo* protein targeting experiments, mass spectrometry-based analyses of organellar fractions, protein-protein interaction data and bioinformatics tools for subcellular localization prediction. Based on the latter approach, purity of our mitochondrial fractions can be estimated to be in the range of 90% (87 to 94%, Supplemental Fig. S2). Traces of chloroplasts were present in our fractions. However, even Rubisco, which is considered to be the most abundant chloroplast protein, was detectable only as a very faint spot on the 2D BN/SDS gel of the L-mito fraction (Fig. 1). Finally, quantitative lipid analysis revealed the purity of our mitochondrial fraction. MGDG, DGDG, PG and SQDG are the main components of plastidial membranes (Hölzl and Dörmann, 2019). In *Arabidopsis* chloroplasts, the ratio of MGDG : DGDG : PG : SQDG is about 1:0.5:0.1:0.3 (Awai et al., 2006). In contrast, a distinct ratio of 1:0.3:0.4:0 was detected in L-mito in this study (Fig. 2). Moreover, only a few specific MGDG and DGDG molecular species are enriched in L-mito compared to L-TE (Fig. 3B), suggesting that the cross-contamination from intact plastids or bulk plastidial membranes are insignificant. Interestingly a similar observation was recently made when tagged mitochondria were isolated by affinity purification (Niehaus et al., 2020). However, in previous studies on mitochondria isolated from cell cultures the amount of MGDG was much lower (~1.4 mol % vs. ~15 mol % in Fig. 2) than in the mitochondrial fraction analyzed here from leaves (Jouhet et al., 2004; Michaud et al., 2016). This can only be explained by a higher number of membrane contacts between plastids and mitochondria in leaves as has been observed before in case of phosphate starved cell cultures (Jouhet et al., 2004). On the other hand, more than 85 % of the subunits of the MTL complex (deduced from its composition in yeast mitochondria, Fig. 7) and many OE-localized proteins (Fig. 8) were identified in the mitochondrial samples via our proteomics approach, strongly suggesting that a close contact between mitochondria, the ER and chloroplasts exists in our preparations as has been observed before in *Arabidopsis* cell cultures (Jouhet et al., 2004; Michaud et al., 2016). We conclude that the identified chloroplast lipids and proteins rather originate from small pieces of plastidial membranes and, importantly, the mitochondria-plastid contact sites, which are present in our mitochondrial fraction and contribute to the measured lipid composition. While the protein complexes connecting ER and mitochondria have been well described in yeast, their roles in mediating and/or facilitating lipid translocation are less defined in plants. In contrast to yeast, to our knowledge only one tethering protein has been described in flowering plants up to now (Michaud et al., 2016).

With the proteomic and lipidomic datasets as well as the online resources in hand, we assigned central lipid biosynthesis reactions and refined models for a possible exchange of lipid molecules between mitochondria, ER and plastids in plants in Fig. 9. Previous studies in yeast and mammalian cells have shown that mitochondria are capable of synthesizing PE, PA, PG and CL (Flis and Daum, 2013; Horvath and Daum, 2013; Tatsuta et al., 2014). Here, we expand the knowledge that at least the glycerolipids PE and CL can be generated and/or modified in Arabidopsis leaf mitochondria. Considering PE biosynthesis, the rate-limiting enzymes in CDP-ethanolamine and PS decarboxylation pathways, PECT1 and PSD1, respectively, were identified in our mitochondrial samples with higher abundance in C-mito (Fig. 4). It suggests that the generation of mitochondrial PE in the cell culture may be more active in comparison to the leaves. Moreover our observation was supported by earlier studies that localized PECT1 at the mitochondrial periphery and PSD1 in mitochondria in *Arabidopsis* (Mizoi et al., 2006; Nerlich et al., 2007). PE is one of the most abundant glycerophospholipids in biological membranes. Therefore, a high demand is expected to supply mitochondria for their biogenesis in actively dividing cell cultures. The other major component of biological membranes is PC. It is considered to be synthesized in the ER and then transferred through protein complexes connecting ER and mitochondria to mitochondria. The structural information of the enriched lipid species provides evidences for the biosynthesis in the ER as well. Many PE and PC species in mitochondrial samples have longer acyl moieties with 22 to 26 carbons, which can only be synthesized in the ER (Haslam and Kunst, 2013). Conventionally, the majority of PS biosynthesis was assumed to take place in the ER and before being transferred to mitochondria as well. PS and PE, although they are able to interconvert, have distinct lipid profiles from each other, suggesting that only selected lipid species are the substrates of PSD1. In addition, inositol polyphosphate phosphatase 1 (SAL1), which is an enzyme that is involved in phosphoinositide degradation and generates inositol phosphates, was identified in our mitochondrial samples with higher abundance in C-mito comparing to L-mito. This finding confirmed a previous report that localized it in both mitochondria and plastids (Estavillo et al., 2011). Inositol phosphates play crucial roles in many biological processes including gene expression and regulation of cell death through sphingolipids (Alcázar-Román and Wente, 2008; Donahue et al., 2010). Removal of head groups from glycerophospholipids, mostly PC and PE, by phospholipase Dα1 (PLDα1) results in PA, serving as important precursor in other glycerophospholipid biosynthesis pathways (Fig. 9). Comparing to L-TE, the most enriched PA species in L-mito carry 16:0 and 18:0 fatty acyl moieties. This corresponds to the enriched PI, PG and DAG species in L-mito, suggesting that PA may serve as an interconverting hinge between these glycerolipids in plant mitochondria. While our study detected PLDα1 preferentially in C-mito samples, previous studies localized it in the cytosol or at the plasmamembrane (Du et al., 2013). This may suggest that PLDα1 localizes to contact sites either between plasmamembrane and mitochondria or between ER and mitochondria. In addition, two glycerol-3-phosphate dehydrogenases were identified (SDP6 and GPDHp). Whereas SDP6 was detected in mitochondria before, GPDHp was localized in the plastid (Nandi et al., 2004; Quettier et al., 2008).

**Figure 9.**
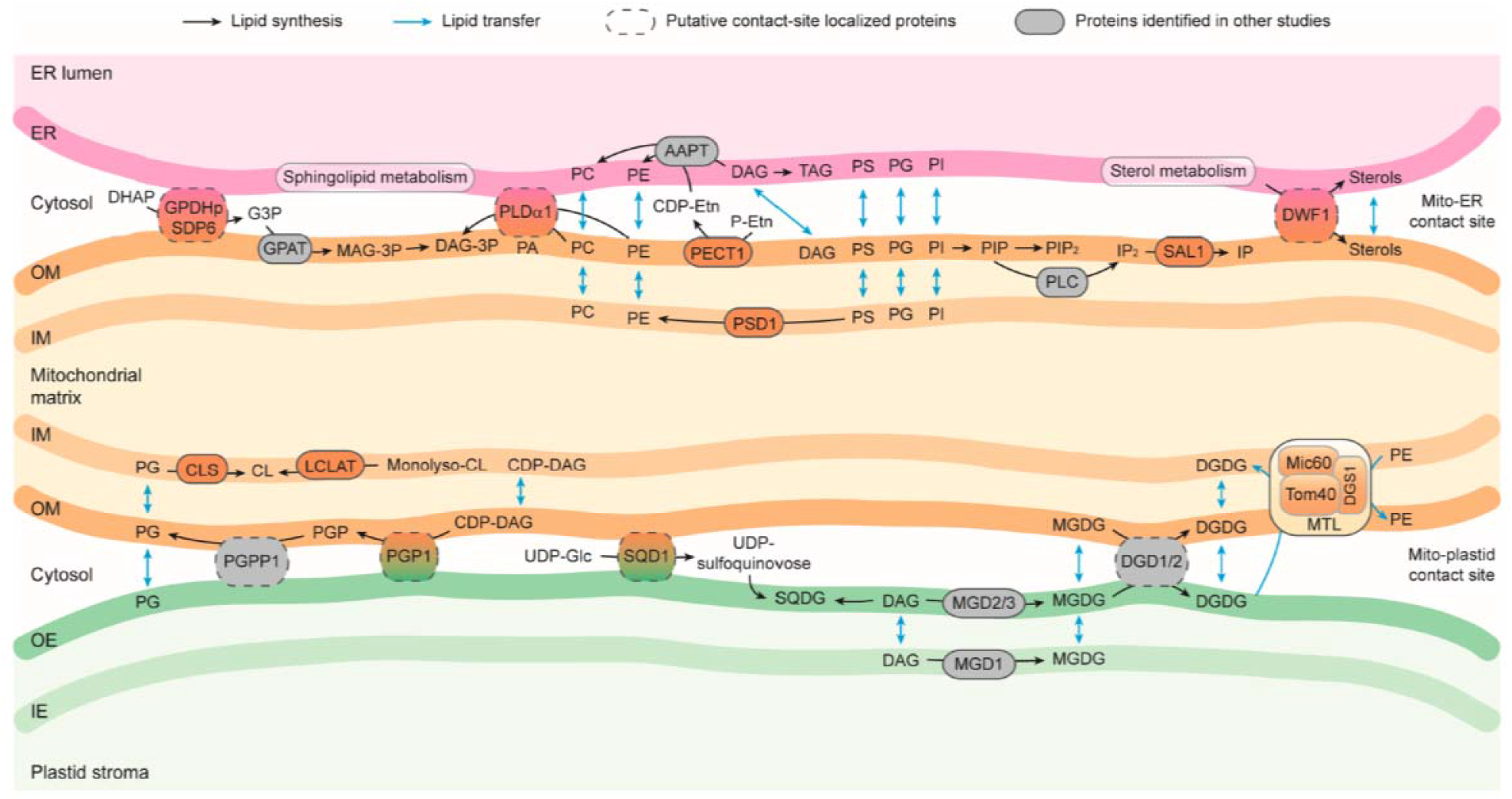
Model of lipid biosynthesis and trafficking within and between mitochondria, ER and plastids in *Arabidopsis.* Lipid synthesis and transfer between membranes are indicated by black and blue arrows, respectively. Proteins identified in L-mito or C-mito with additional ER or plastidic localization based on The Arabidopsis Information Resource (TAIR; www.arabidopsis.org) and the Subcellular localization database for Arabidopsis proteins (SUBAcon; www.suba.live) are considered as putative contact-site localized proteins (dashed frame). Lipid biosynthesis-related proteins indicated in other studies are depicted in grey. Full names and functions of involved proteins are itemized in Supplemental Tab. S1. OM, mitochondrial outer membrane; IM, mitochondrial inner membrane; OE, plastid outer envelope; IE, plastid inner envelope; MTL, mitochondrial transmembrane lipoprotein complex. DHAP, dihydroxyacetone phosphate; G3P, glyceraldehyde 3-phosphate; MAG, monoacylglycerol; DAG, diacylglycerol; Etn, ethanolamine; TAG, triacylglycerol; IP, inositol phosphate; PGP, phosphatidylglycerol phosphate; SQDG, sulfoquinovosyl diacylglycerol.

An enzyme being involved in synthesizing PG from PA is PGP1. It was found in our proteomic analysis in C-mito samples (Fig. 4). CL can be synthesized through condensation of PG and DAG by CLS, or addition of acyl chains to monolyso-CL by monolysocardiolipin acyltransferase (LCLAT or Tafazzin). All identified enzymes involving in PG and CL biosynthesis, PGP1, CLS and LCLAT, are more abundant in C-mito compared to L-mito. PGP1 was previously found to localize to both mitochondria and plastids (Babiychuk et al., 2003), while CLS was localized to mitochondria (Katayama et al., 2004). This suggests an intensive interaction between mitochondria and plastids in cell cultures because (1) high amounts of proteins localized in contact sites imply a closer connection between the two organelles; (2) more CLS is required to convert the plastid-derived PG which is transferred to mitochondria presumably via membrane contact sites; (3) active energy generation is vital for actively dividing cells and thus a high amount of CL is required to assemble the respiratory chain complexes in mitochondria. This is further supported by the specific PG species found in L-TE. L-TE contains substantial amounts of 16:1 (probably *trans*-Δ^3^-16:1)/18:X PG species (~35 mol %) (Fig. 3), which are molecular species synthesized entirely within plastids and probably exclusively localized there (Hölzl and Dörmann, 2019). Therefore, it is very likely that the high amounts of these PG species in L-TE can be attributed to the abundance of plastid membranes particularly thylakoids. Interestingly, the L-mito fraction also contained significant levels (~8 mol %) of 16:1/18:X PG species. These PG species may be not in mitochondria but be derived from the mitochondrion-plastid contact sites or contaminations of plastid membranes (Fig. 9). Nevertheless PG which is directly after its uptake converted into CL in mitochondria (Figs. 2 and 9).

Typically, glyceroglycolipids such as MGDG, DGDG and SQDG are considered to be synthesized in plastids and are transferred to other organelles upon stress. For instance, DGDG is transferred from plastidial membranes to other compartments including mitochondrial membranes and the plasma membrane during phosphate starvation to compensate the loss of PC and PE (Jouhet et al., 2004). In addition to vesicular transportation and lipid trafficking via contact sites, emerging evidences support the hypothesis of lipid synthesis in *trans* in yeast and plants (Mehrshahi et al., 2013; Tavassoli et al., 2013; Michaud et al., 2017). That is, enzymes located at one membrane might be capable of catalyzing a reaction on another membrane when in close proximity. In this way, neither tethering proteins nor massive lipid remolding under stress conditions is required, if the mitochondria can acquire lipids without setting up the junctions to other organelles. Therefore, we suspect that the DGDG biosynthesis enzymes, DGD1/2, catalyze the reactions in *trans* or at the mitochondria-plastid contact sites, providing DGDG molecules to compensate the loss of glycerophospholipids in real time (Fig. 4b). In addition, several subunits of the MTL complex were identified in our proteomic approach including Tom40, Mic60, DGS1 etc. (Fig. 7). It establishes a lipid trafficking system removing DGDG from plastids and allocating PE between the IM and OM. Rapid remodeling of the mitochondrial membranes is supported by in *trans* lipid biosynthesis and the MTL complex, likely in close cooperation with other complexes. However, the underlying mechanism of this lipid transportation machinery is still largely unknown. SQDG synthase (SQD1) was identified exclusively in C-mito, but no SQDG molecules were detected (Fig. 4b). Earlier reports localized SQD1 in chloroplasts (Essigmann et al., 1998), suggesting that SQD1 is either co-purified with plastid envelope membranes from cell cultures or is localized in .

Sphingolipid biosynthesis takes place at the ER membrane and subsequently in the Golgi apparatus for the addition of carbohydrate residues. Interestingly, in the last decade, several sphingolipid-metabolizing enzymes have been identified in mitochondria purified from yeast and mammalian systems, including Cer synthase and ceramidase (Bionda et al., 2004; Kitagaki et al., 2007; Novgorodov et al., 2011). In animal models, mitochondria-synthesized Cer plays a crucial role in cerebral ischemia-induced mitochondrial dysfunction. The connection between sphingolipids and mitochondria-promoted apoptosis has been proposed in plants as well. Recent studies in *Arabidopsis* have demonstrated that the ratio between LCB-P and Cer is involved in maintaining the balance of cell survival and apoptosis (Watanabe et al., 2013; Bi et al., 2014). However, no sphingolipid biosynthesis enzyme has been identified in plant mitochondria hereto. GIPCs are the most abundant sphingolipids in mitochondria, although still minor comparing to glycerophospholipids. It is known that PM-localized GIPCs are important in signal transduction and intercellular recognition (Lenarčič et al., 2017; Ali et al., 2018). GIPCs may as well establish the communication between mitochondria and other organelles, although further analysis is required to understand the functions of sphingolipids in plant mitochondria.

Sterols, although taking part in both biotic and abiotic stress responses, are at low abundance in mitochondrial samples. Sterol biosynthesis primarily takes place in the ER (Schaller, 2003). Among them, sterol C-24 reductase (DWF1) was identified both in L-mito and C-mito, with higher abundance in C-mito. DWF1 has been proposed to mediate the biosynthesis of all phytosterols with higher specificity towards campesterol in Arabidopsis, but this protein was localized to ER membranes (Klahre et al., 1998; Youn et al., 2018). Therefore, we suggest DWF1 as an additional contact site localized protein (Fig. 7). High amount of campesterol was detected in L-mito, suggesting the existence of an onsite biosynthesis and/or sterol transporter to facilitate the import of campesterol from the ER. However, unlike mammalian cells wherein cholesterol transport proteins such as steroidogenic acute regulatory protein, StAR (Clark et al., 1995), and MLN64 (Charman et al., 2010) have been identified, little is known about sterol transporters in plants.

## Conclusions

In summary, we expanded the knowledge regarding lipid biosynthesis and modification in plant mitochondria by performing a global lipidome analysis of Arabidopsis leaf mitochondria to provide their in-depth lipid molecular species profile including glycerolipids, sphingolipids and sterols. Our proteome data confirmed that PE and CL can be synthesized in these organelles partially by the assistance of putative contact-site localized proteins and / or in *trans* lipid biosynthesis. Based on the proteomic results, we propose and confirm the existence of membrane contact site-localized proteins and their aspects in lipid biosynthesis pathways. This study serves as a foundation for additional researches in unveiling the functional roles of mitochondrial lipids and the mechanisms of mitochondria-dependent signaling pathways.

## Materials and methods

### Plant materials and growth conditions

Rosette leaves of wild-type *Arabidopsis thaliana* (L.) Heynh Columbia-0 were used for both extracting total lipid extract and purifying mitochondria. After sowing the seeds in pots with three-day cold stratification at 4 °C, seedlings were grown under 16 h-day length at 24 °C with 60 % relative humidity and 150 μmol photons m^-2^ sec^-1^ for one week. Young seedlings were transferred to large trays and grown with equal spacing for another three weeks before further experimental procedures.

### Suspension culture of *Arabidopsis thaliana*

Suspension cultures were established starting from sterilized seeds of *Arabidopsis thaliana* wild-type Columbia-0, grown on MS medium plates containing 0.8 % agar. Plant pieces were transferred to B5 medium agar plates and cultivated for several weeks in the dark for callus induction. Callus was finally transferred into liquid B5 medium including 3 % (w/v) sucrose, 0.01 % (w/v) 2,4-dichlorophenoxyacetic acid and 0.001 % (w/v) kinetin. Cultures were incubated on a shaker at 24 °C in the dark. Callus was transferred weekly into new liquid medium (3 g / 100 ml).

### Mitochondria isolation

#### From Arabidopsis rosettes

The mitochondria isolation procedure was described previously (Schikowsky et al., 2018). About 200 g of four-week-old *Arabidopsis* rosettes were collected and homogenized at 4 °C with a waring blender in 1 liter disruption buffer (0.3 M sucrose, 60 mM TES, 25 mM tetrasodium pyrophosphate, 10 mM potassium dihydrogen phosphate, 2 mM EDTA, 1 mM glycine, 1 % PVP40, 1 % BSA, 50 mM sodium ascorbate, 20 mM cysteine; pH 8.0) by three times for 10 sec with 30 sec intervals. The following procedures were performed on ice or at 4 °C. Two layers of miracloth with supporting gauze were used to filter the homogenate into a beaker. The remaining plant debris was first grinded with additional sea sand for 10 min by mortar and pestle, and then filtered again through miracloth. The filtrates were combined and centrifuged at 2,500 g for 5 min to eliminate the cell debris. Centrifugation with higher speed at 15,250 g for 15 min was applied on the supernatant to pellet mitochondria and other organelles. The resulting pellets were resuspended with a paintbrush in wash buffer (0.3 M sucrose, 10 mM TES, 10 mM potassium dihydrogen phosphate; pH 7.5). The samples were adjusted to the final volume of 12 ml with wash buffer and transferred to a Dounce homogenizer. Two strokes of pestle were performed to disrupt large organelles like chloroplasts. Aliquots of 1 ml samples were transferred carefully to Percoll gradients which had been established beforehand by 69,400 g centrifugation for 40 min in one-to-one ratio of Percoll and Percoll medium (0.6 M sucrose, 20 mM TES, 2 mM EDTA, 20 mM potassium dihydrogen phosphate, 2 mM glycine; pH 7.5). Mitochondria were separated from other components by centrifuging in the gradients at 17,400 g for 20 min. The resulting mitochondrial fractions formed white clouds at the bottom half of the gradients and were collected by Pasteur pipettes to clean centrifuge tubes. The clean-up procedures were performed three to five times by filling up wash buffer in the centrifuge tubes and pelleting the mitochondria with 17,200 g for 20 min, until the resulting pellet was firm. After each washing steps, two to three pellets were combined in one tube until all mitochondria from one biological replicate were pooled together. The mitochondrial pellets were weighed, resuspended with wash buffer and aliquoted at the concentration of 0.1 g/ml.

#### From cell cultures

Mitochondria isolation from *Arabidopsis thaliana* suspension cell culture was carried out as described before (Farhat et al., 2019). About 200 g fresh cells were harvested and homogenized using disruption buffer (450 mM sucrose, 15 mM MOPS, 1.5 mM EGTA, 0.6 % (w/v) PVP40, 2 % (w/v) BSA, 10 mM sodium ascorbate, 10 mM cysteine; pH 7.4) and a waring blender. During several washing steps, cell fragments were removed (centrifugation twice for 5 minutes at 2,700 g and once for 5 minutes at 8,300 g). Crude mitochondria were pelleted at 17,000 g for 10 minutes, resuspended in washing buffer (0.3 M sucrose, 10 mM MOPS, 1 mM EGTA; pH 7.2), homogenized using a Dounce homogenisator and loaded onto discontinuous Percoll gradients (phases of 18 %, 23 % and 40 % Percoll in gradient buffer (0.3 M sucrose, 10 mM MOPS, 0.2 mM EGTA; pH 7.2)). After ultracentrifugation (90 minutes, 70,000 g), purified mitochondria were collected from the 23 %-40 % interphase. For Percoll removal, several washing steps (10 minutes, 14,500 g) were performed using resuspension buffer (0.4 M mannitol, 1 mM EGTA, 10 mM tricine; pH 7.2) to gain a firm pellet of purified mitochondria.

Each of the three independently purified mitochondria populations from 4-week old rosette leaves (L-mito) and 200 g cell cultures (C-mito), respectively were used for all experiments.

### BN/SDS-PAGE

Gel electrophoresis procedures (blue-native (BN) and SDS PAGE) were performed as described previously (Senkler et al., 2018), based on the published protocol given in (Wittig et al., 2006).

### Label-free quantitative shotgun proteomics

#### Protein sample preparation

Sample preparation for shotgun proteome analysis was performed as described before (Thal et al., 2018). The protein content of the mitochondrial fractions was determined using a Bradford assay kit (Thermo Scientific, Rockford, USA). 50 μg protein of each sample were loaded onto a SDS gel for sample purification (Thal et al., 2018). Electrophoresis was stopped when the proteins reached the border between the stacking and the separating gel. Gels were subsequently incubated in fixation solution (15% (v.v) ethanol; 10% (v.v) acetic acid) for 30 min, stained for 1 h with Coomassie Brilliant Blue G250, and finally the protein band at the border of the two gel phases was cut out into cubes with edge lengths of approximately 1 mm. Trypsination of the proteins was carried out as described previously (Fromm et al., 2016).

#### Shotgun proteomic LC-MS analysis

Label-free quantitative mass spectrometric analyses of whole mitochondrial protein samples from cell culture and long day *Arabidopsis thaliana* leaves were performed as outlined before (Thal et al., 2018) using an Ultimate 3000 UPLC coupled to a Q Exactive Orbitrap mass spectrometer (Thermo Scientific, Dreieich, Germany).

#### Data processing

In a first step, the resulting MS data were processed using the Proteome Discoverer Software (Thermo Fisher Scientific, Dreieich, Germany) and searched with the Mascot search engine (www.matrixscience.com) against the tair10 protein database (downloaded from www.arabidopsis.org). For quantitative analyses, MS data were further processed as outlined before (Rugen et al., 2019) using the MaxQuant software package (version 1.6.4.0), the Andromeda search engine (Cox and Mann, 2008) and the tair10 protein database. For determination of sample purity, peptide intensities were used, combined with the subcellular locations of assigned proteins as given by SUBAcon from the SUBA platform (www.suba.live) (Hooper et al., 2017). A proteomic heatmap was generated using the NOVA software (www.bioinformatik.uni-frankfurt.de/tools/nova/index.php)(Giese et al., 2015). Identified proteins of all six datasets (three biological replicates of leaf total mitochondrial protein (L-mito 1,2,3) and cell culture total mitochondrial protein (C-mito 1,2,3)) were hierarchically clustered in a heatmap based on iBAQ (<u>i</u>ntensity <u>B</u>ased_<u>A</u>bsolute <u>Q</u>uantification) values (for primary results see Supplemental Tab. S1).

### Lipid extraction

Total lipid extract from leaf (L-TE) was obtained as described (Tarazona et al., 2015). Briefly, mixture of 150 mg frozen leaf powder or 100 mg mitochondria and 6 ml extraction buffer (propan-2-ol : hexane : water, 60:26:14 (v.v.v)) was incubated at 60 °C with shaking for 30 min. After centrifugation at 800 g for 20 min, the clear supernatant was transferred to clean tubes and evaporated under stream of nitrogen gas until dryness. Samples were reconstituted in 800 μl of TMW (tetrahydrofuran (THF) : methanol : water, 4:4:1 (v.v.v)). Procedure for extracting the mitochondrial lipids was adjusted by substituting the water fraction of the extraction buffer by equal volume of the mitochondrial aliquot. Further process was continued as described above.

### Lipid quantification

#### TLC separation of lipid classes

Lipid extracts from 500 mg mitochondria and 50 mg leaves were spotted onto TLC 60 plates (20 × 20 cm^2^, Merck KGaA, Darmstadt, Germany) in parallel with the following standards: PC (Larodan, Solna, Sweden), PE, CL, and PG (Avanti Polar Lipids, Birmingham, AL, USA), MGDG, and DGDG (purified from plant lipid extracts). Extracts were developed by a solvent mixture of chloroform : methanol : acetic acid (65:25:8 (v.v.v)). After visualizing the lipid spots under 528 nm UV light, the bands were scrapped out and converted to fatty acid methyl esters (FAME) before GC/FID analysis.

#### Acidic transesterification

Glycerolipids were transesterified by acidic methanolysis (Miquel and Browse, 1992) and converted to fatty acid methyl esters (FAME). The lipid-bound silica powder from corresponding spots on the TLC plate were scrapped out and added to 1 ml FAME solution (methanol : toluene : sulfuric acid : dimethoxypropane, 33:17:1.4:1 (v.v.v.v)) with 5 μg tripentadecanoin as an internal standard. After 1 h incubation at 80 °C, 1.5 ml saturated NaCl solution and 1.2 ml hexane were added subsequently. The resulting FAME was collected from the hexane phase, dried, and reconstituted in 10 μl acetonitrile.

#### GC/FID analysis

Lipid-bound fatty acids were analyzed after converting to FAMEs by a 6890N Network GC/FID System with a medium polar cyanopropyl DB-23 column (30 m × 250 μm × 25 nm; Agilent Technologies, Waldbronn, Germany) using helium as the carrier gas at 1 ml min^-1^. Samples were injected at 220 °C with an Agilent 7683 Series injector in split mode. After 1 min at 150 °C, the oven temperature raised to 200 °C at the rate of 8 °C min^-1^, increased to 250 °C in 2 min, and held at 250 °C for 6 min. Peak integration was performed using the GC ChemStation (Agilent Technologies).

### Lipid derivatization

Phosphate-containing lipids - phosphatidic acids (PA), phosphoinositides (PIPs) and long-chain base phosphates (LCB-P) - were derivatized to enhance their chromatographic separation and mass spectrometric detection. Methylation procedure was applied on PA and PIPs as followed. Aliquots of 100 μl lipid extracts were first brought to dryness under stream of nitrogen gas and reconstituted with 200 μl methanol. Methylation reaction took place after the supply of 3.3 μl trimethylsilyldiazomethane. After 30 min incubation at room temperature, the reaction was terminated by neutralizing with 1 μl of 1.7 M acetic acid. Samples were dried under nitrogen gas and redissolved in 100 μl TMW. Acetylation procedure was applied on LCB-P. Aliquots of 100 μl lipid extracts were brought to dryness and reconstituted with 100 μl pyridine and 50 μl acetic anhydride. After 30 min incubation at 50 °C, samples were dried under stream of nitrogen gas with 50 °C water bath. To redissolve the samples, 100 μl TMW was used as the final solvent to proceed with lipid analysis.

### Global lipidomic analysis with LC-MS

Analysis conditions and system setup were as described (Tarazona et al., 2015). Samples were separated by an ACQUITY UPLC system (Waters Crop., Milford, MA, USA) with a HSS T3 column (100 mm x 1 mm, 1.8 μl; Waters Crop.), ionized by a chip-based nanoelectrospray using TriVersa Nanomate (Advion BioScience, Ithaca, NY, USA) and analyzed by a 6500 QTRAP tandem mass spectrometer (AB Sciex, Framingham, MA, USA). Aliquots of 2 μl were injected and separated with a flow rate of 0.1 ml min^-1^. The solvent system composed of methanol : 20 mM ammonium acetate (3:7 (v.v)) with 0.1 % (v.v) acetic acid (solvent A) and THF : methanol : 20 mM ammonium acetate (6:3:1 (v.v.v)) with 0.1 % (v.v) acetic acid (solvent B). According to the lipid classes, different linear gradients were applied: start from 40 %, 65 %, 80 % or 90 % B for 2 min; increase to 100 % B in 8 min; hold for 2 min and re-equilibrate to the initial conditions in 4 min. Starting condition of 40 % solvent B were utilized for long-chain bases (LCB) and phosphorylated long-chain bases (LCB-P); 80 % for diacylglycerol (DAG); 90 % for steryl esters (SE); 65 % for the remaining lipid classes. Retention time alignment and peak integration were performed with MultiQuant (AB Sciex). Quantitative results were calculated according to the amount of internal standards.

### Biosynthesis pathways construction

The Arabidopsis Information Resource (TAIR; www.arabidopsis.org), the Subcellular localization database for Arabidopsis proteins (SUBAcon; www.suba.live), Kyoto Encyclopedia of Genes and Genomes (KEGG; www.genome.jp/kegg) and the shotgun proteomic analyses of L-mito and C-mito were combined to construct the lipid biosynthesis pathways in plant mitochondria. Additionally, lipidomics data are depicted in the pathways to illustrate the biosynthetic fluxes.

## Supplemental Data

**Supplemental Figure S1.** Purity inspection of mitochondrial protein complexes and supercomplexes by two-dimensional blue-native/SDS PAGE. Mitochondrial fractions isolated from Arabidopsis leaves (L-mito 1 and L-mito 3) and from Arabidopsis cell cultures (C-mito1 and C-mito 3) were separated by 2D PAGE and Coomassie-stained (corresponding gels of fractions L-mito 2 and C-mito 2 see Fig. 1). Numbers on top and to the left of the 2D gels refer to the masses of standard protein complexes / proteins (in kDa), the roman numbers above the gels to the identity of OXPHOS complexes (see Fig. 1 for detailed information). The arrows indicate the large (L; 53,5 kDa) and the small (S; 14.5 kDa) subunit of Rubisco. To further confirm the reproducibility of the preparations, 2D blue-native/SDS reference gels for mitochondrial and chloroplast fractions (Mito-ref, Cp-ref) from *A. thaliana* are given to the bottom of the figure (gels were taken from (Klodmann et al., 2011) and (Behrens et al., 2013)). Identity of the protein complexes visible on the chloroplast reference gel: PSI – photosystem I; PSII – photosystem II; Rub – Rubisco; LHCII – light harvesting complex II.

**Supplemental Figure S2.** Purity of mitochondrial fractions as determined by label-free quantitative shot gun proteomics. Three mitochondrial fractions isolated from Arabidopsis leaves (L-mitos) and cell cultures (C-mitos) were analysed. Summed-up peptide intensities were calculated for subcellular compartments based on protein assignments as given by SUBAcon (The Subcellular localization database for Arabidopsis proteins; www.suba.live). Blue: mitochondria; green: plastids; gray: others; numbers in %.

**Supplemental Figure S3.** Fatty acid profiles of the glycerolipids from L-mito, L-TE, C-mito and C-TE. Heat maps illustrate the difference of the fatty acid distribution based on LC-MS/MS analyses. The heat map colors represent mean intensity values according to the color map on the low right-hand side. Each column represents one lipid class wherein the acyl moieties of all species are listed and summarized to 100 %. Data represent mean values in mol % from three independent experiments which were also used for proteomics (Supplemental Tab. S1).

**Supplemental Figure S4.** Workflow for the construction of the biosynthesis pathways. Multiple databases were combined to build the lipid biosynthesis pathways in mitochondria, The Arabidopsis Information Resource (TAIR; www.arabidopsis.org), the Subcellular localization database for Arabidopsis proteins (SUBAcon; www.suba.live), Kyoto Encyclopedia of Genes and Genomes (KEGG; www.genome.jp/kegg) and the proteomic datasets of isolated mitochondria in this study. Information of protein localizations and backbones of the biosynthesis pathways were obtained from TAIR, SUBAcon and KEGG, respectively.

**Supplemental Figure S5.** Additional lipid biosynthesis enzymes which do not match with analyzed lipid classes in this study. Enzymes of (a) fatty acid biosynthesis, (b) unsaturated fatty acid biosynthesis, (c) lipoic acid metabolism and (d) ubiquinone and other terpenoid-quinone biosynthesis illustrate the general lipid biosynthesis in Arabidopsis mitochondria. Heat maps visualize the protein abundance based on shotgun proteomic analyses of three independent experiments of purified mitochondria from leaves and cell culture. The heat map colors represent mean intensity values according to the color map on the low right-hand side. Blue: proteins identified in the proteomic analysis of this study and also predicted to localize in mitochondria; green: proteins identified in the proteomic analysis of this study but predicted to localize in other organelles; bold font: exclusively localized in mitochondria; italic font: only identified in one of the mitochondrial populations. Further full names and functions are itemized in Supplemental Tab. S3. In the heat maps, proteins with high and low abundance are depicted in yellow and black, respectively. Predicted protein localization was based on The Arabidopsis Information Resource (TAIR; www.arabidopsis.org) and the Subcellular localization database for Arabidopsis proteins (SUBAcon; www.suba.live).

**Supplemental Table S1.** Overall proteins identified in the proteomic approach.

**Supplemental Table S2.** Overall lipids identified in the lipidomic approach.

**Supplemental Table S3.** Lipid biosynthesis-related proteins identified in the proteomic approach.

<u>**Supplemental Table S4.** List of subunits of the MTL identified in the proteomic approach.</u>

<u>**Supplemental Table S5.** Proteins assigned to plastids regarding to SUBAcon.</u>

## Acknowledgements

We are grateful for the technical assistance from Sabine Freitag. YL has been a doctoral student of the Ph.D. program “Molecular Biology” – International Max Planck Research School and the Göttingen Graduate School for Neurosciences, Biophysics, and Molecular Biosciences (GGNB) (DFG grant GSC 226) at the Georg August University Göttingen.

