## Supplementary material for "Defining the lipidome of Arabidopsis leaf mitochondria: Specific lipid complement and lipid biosynthesis capacity": Supplmental Figures S1-S5

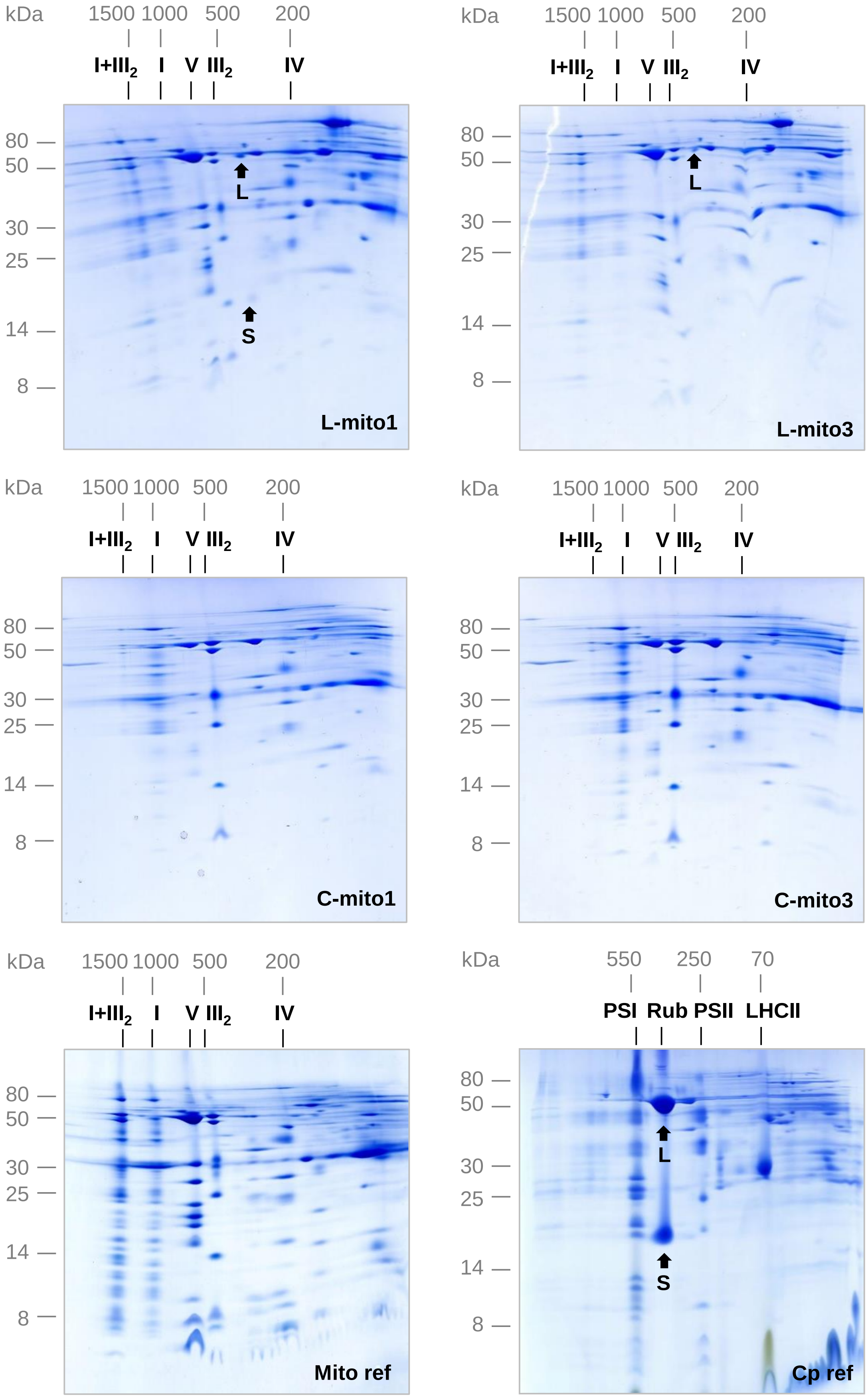


**Supplemental Figure S1.** Purity inspection of mitochondrial protein complexes and supercomplexes by two-dimensional blue-native/SDS PAGE. Mitochondrial fractions isolated from Arabidopsis leaves (L-mito 1 and L-mito 3) and from Arabidopsis cell cultures (C-mito1 and C-mito 3) were separated by 2D PAGE and Coomassie-stained (corresponding gels of fractions L-mito 2 and C-mito 2 see Fig. 1). Numbers on top and to the left of the 2D gels refer to the masses of standard protein complexes / proteins (in kDa), the roman numbers above the gels to the identity of OXPHOS complexes (see Fig. 1 for detailed information). The arrows indicate the large (L; 53,5 kDa) and the small (S; 14.5 kDa) subunit of Rubisco. To further confirm the reproducibility of the preparations, 2D blue-native/SDS reference gels for mitochondrial and chloroplast fractions (Mito-ref, Cp-ref) from A. thaliana are given to the bottom of the figure (gels were taken from Klodmann et al., 2011 and Behrens et al., 2013). Identity of the protein complexes visible on the chloroplast reference gel: PSI – photosystem I; PSII – photosystem II; Rub – Rubisco; LHCII – light harvesting complex II.

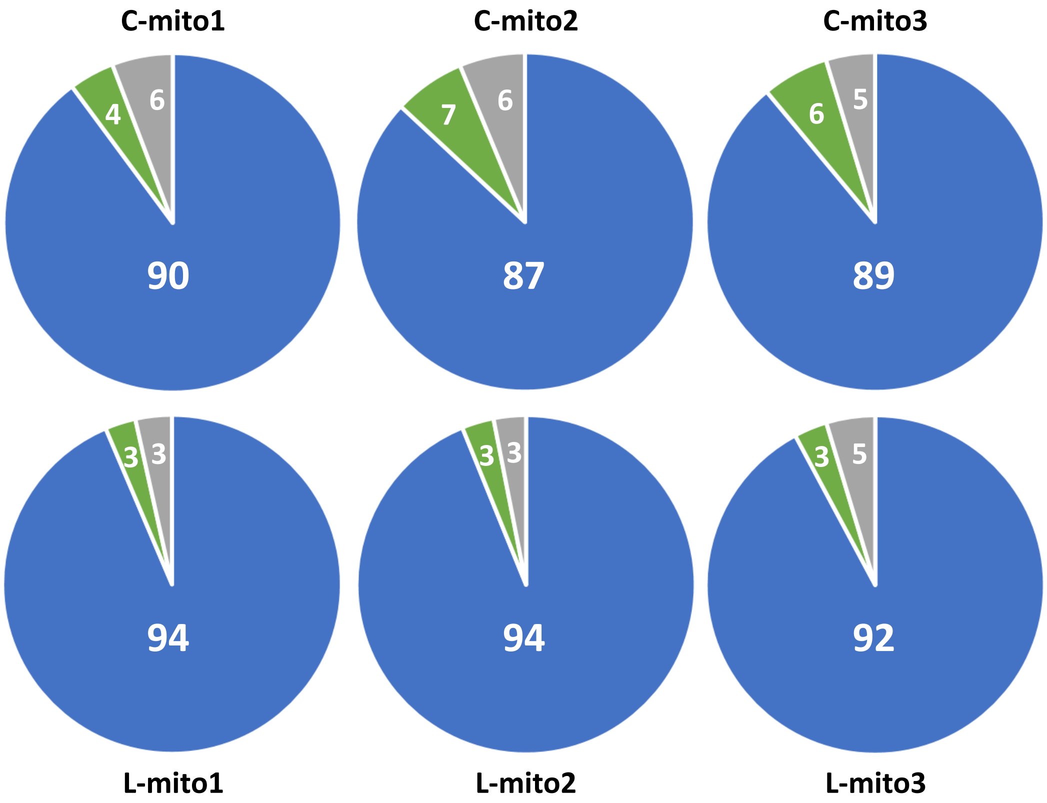


**Supplemental Figure S2.** Purity of mitochondrial fractions as determined by label-free quantitative shot gun proteomics. Three mitochondrial fractions isolated from Arabidopsis leaves (L-mitos) and cell cultures (C-mitos) were analysed. Summed-up peptide intensities were calculated for subcellular compartments based on protein assignments as given by SUBAcon (The Subcellular localization database for Arabidopsis proteins; www.suba.live**).** Blue: mitochondria; green: plastids; gray: others; numbers in %.

**L-mito**

| Mol % | PC | PE | PS | PA | PI | PIP | PIP2 | PG | CL | MGDG | DGDG | DAG |
|---|---|---|---|---|---|---|---|---|---|---|---|---|
| 16:0 | 12.6 | 7.2 | 10.5 | 24.5 | 43.3 | | 45.4 | 44.1 | 0.3 | 2.5 | 8.6 | 21.6 |
| 16:1 | 0.7 | 0.4 | 0.5 | 0.8 | 0.6 | | | 4.9 | 0.4 | 1.3 | 0.5 | 1.2 |
| 16:2 | | | | | | | | | | 2.1 | 0.9 | |
| 16:3 | | | | | | | | 0.0 | | 26.8 | 7.3 | |
| 18:0 | 1.0 | 0.4 | 28.8 | 2.6 | 2.5 | | | 1.1 | 1.8 | 0.4 | 0.8 | 2.9 |
| 18:1 | 4.7 | 2.4 | 2.0 | 5.2 | 2.0 | | 6.4 | 5.1 | 6.5 | 2.0 | 1.3 | 5.1 |
| 18:2 | 41.8 | 51.2 | 33.1 | 36.0 | 28.3 | | 30.9 | 22.5 | 23.9 | 5.6 | 4.4 | 32.2 |
| 18:3 | 39.0 | 37.9 | 13.4 | 30.9 | 23.3 | | 17.3 | 22.3 | 67.2 | 59.3 | 76.2 | 34.5 |
| 20:0 | 0.0 | 0.1 | | | | | | | | | | 0.4 |
| 20:1 | 0.1 | 0.1 | | | | | | | | | | 0.3 |
| 20:2 | 0.2 | 0.2 | | | | | | | | | | |
| 22:0 | 0.0 | 0.1 | | | | | | | | | | 0.5 |
| 22:1 | 0.0 | | | | | | | | | | | 0.4 |
| 24:0 | | 0.1 | 4.2 | | | | | | | | | 0.6 |
| 24:1 | | 0.1 | 5.4 | | | | | | | | | 0.4 |
| 26:0 | | | 0.2 | | | | | | | | | |
| 26:1 | | | 1.8 | | | | | | | | | |

**L-TE**

| Mol % | PC | PE | PS | PA | PI | PIP | PIP2 | PG | CL | MGDG | DGDG | DAG |
|---|---|---|---|---|---|---|---|---|---|---|---|---|
| 16:0 | 20.2 | 26.0 | 22.6 | 17.8 | 40.8 | 5.6 | 46.3 | 32.2 | | 1.1 | 9.5 | 26.5 |
| 16:1 | 0.7 | 0.4 | 1.2 | 0.6 | 1.3 | 0.8 | | 18.0 | | 1.7 | 0.4 | 1.1 |
| 16:2 | | | | | | | | | | 2.7 | 1.1 | |
| 16:3 | | | | | | | | 0.5 | | 32.9 | 5.2 | |
| 18:0 | 1.7 | 1.5 | 2.8 | 1.2 | 1.6 | 27.9 | | 0.3 | | 0.1 | 0.7 | 3.8 |
| 18:1 | 4.5 | 3.3 | 1.6 | 3.1 | 1.3 | 11.9 | 3.6 | 3.1 | | 1.4 | 1.2 | 4.6 |
| 18:2 | 36.8 | 40.3 | 22.6 | 38.5 | 21.5 | 30.5 | 34.9 | 11.8 | | 4.1 | 5.1 | 36.0 |
| 18:3 | 35.8 | 27.4 | 30.5 | 38.8 | 33.5 | 23.3 | 15.2 | 34.1 | | 55.9 | 76.8 | 25.8 |
| 20:0 | 0.0 | 0.2 | | | | | | | | | | 0.4 |
| 20:1 | 0.1 | 0.1 | | | | | | | | | | 0.5 |
| 20:2 | 0.1 | 0.2 | | | | | | | | | | |
| 22:0 | 0.0 | 0.2 | | | | | | | | | | 0.4 |
| 22:1 | 0.0 | | | | | | | | | | | 0.4 |
| 24:0 | | 0.2 | 8.4 | | | | | | | | | 0.3 |
| 24:1 | | 0.2 | 8.1 | | | | | | | | | 0.3 |
| 26:0 | | | 1.8 | | | | | | | | | |
| 26:1 | | | 0.4 | | | | | | | | | |

**C-mito**

| Mol % | PC | PE | PS | PA | PI | PIP | PIP2 | PG | CL | MGDG | DGDG | DAG |
|---|---|---|---|---|---|---|---|---|---|---|---|---|
| 16:0 | 11.1 | 8.9 | 17.5 | 20.0 | 43.2 | | 41.3 | 48.0 | 0.5 | 5.0 | 6.9 | 18.0 |
| 16:1 | 1.9 | 1.2 | 0.8 | 1.4 | 1.5 | | | 1.2 | 0.9 | 2.6 | 1.3 | 1.8 |
| 16:2 | | | | | | | | | | 1.4 | 0.4 | |
| 16:3 | | | | | | | | 0.0 | | 10.1 | 1.8 | |
| 18:0 | 0.5 | 0.2 | 17.6 | 1.3 | 1.5 | | | 0.6 | 0.9 | 0.9 | 0.8 | 1.5 |
| 18:1 | 17.4 | 10.7 | 9.8 | 24.1 | 6.4 | | 17.8 | 9.0 | 3.8 | 15.4 | 10.1 | 23.8 |
| 18:2 | 20.5 | 28.7 | 16.0 | 18.0 | 15.5 | | 25.9 | 12.7 | 17.1 | 13.4 | 10.2 | 18.8 |
| 18:3 | 48.5 | 49.9 | 30.0 | 35.1 | 32.0 | | 15.1 | 28.4 | 76.7 | 51.1 | 68.5 | 33.6 |
| 20:0 | 0.0 | 0.0 | | | | | | | | | | 0.6 |
| 20:1 | 0.0 | 0.1 | | | | | | | | | | 0.2 |
| 20:2 | 0.0 | 0.0 | | | | | | | | | | |
| 22:0 | 0.0 | 0.0 | | | | | | | | | | 0.7 |
| 22:1 | 0.0 | | | | | | | | | | | 0.2 |
| 24:0 | | 0.1 | 1.5 | | | | | | | | | 0.4 |
| 24:1 | | 0.1 | 2.0 | | | | | | | | | 0.4 |
| 26:0 | | | 0.2 | | | | | | | | | |
| 26:1 | | | 4.5 | | | | | | | | | |

**C-TE**

| Mol % | PC | PE | PS | PA | PI | PIP | PIP2 | PG | CL | MGDG | DGDG | DAG |
|---|---|---|---|---|---|---|---|---|---|---|---|---|
| 16:0 | 25.7 | 28.7 | 36.5 | 17.9 | 46.2 | | 54.3 | 54.3 | 0.3 | 5.6 | 4.7 | 19.4 |
| 16:1 | 0.2 | 0.9 | | 1.3 | 0.8 | | | 0.8 | 0.6 | 2.1 | 0.6 | 1.2 |
| 16:2 | | | | | | | | | | 0.6 | 0.2 | |
| 16:3 | | | | | | | | | | 9.7 | 4.3 | |
| 18:0 | 1.7 | 2.9 | 3.6 | 1.3 | 0.6 | | | 0.8 | 0.1 | | 0.3 | 1.7 |
| 18:1 | 20.3 | 19.8 | 10.5 | 21.4 | 3.0 | | 11.1 | 7.4 | 1.9 | 17.5 | 5.3 | 22.0 |
| 18:2 | 9.9 | 13.8 | 6.7 | 18.3 | 7.9 | 44.7 | 24.7 | 8.9 | 14.4 | 11.4 | 5.8 | 17.0 |
| 18:3 | 42.2 | 30.6 | 41.8 | 39.8 | 41.6 | 55.3 | 9.9 | 27.8 | 82.8 | 53.0 | 78.9 | 37.3 |
| 20:0 | | 0.9 | | | | | | | | | | 0.4 |
| 20:1 | | 0.3 | | | | | | | | | | 0.1 |
| 20:2 | | 0.0 | | | | | | | | | | |
| 22:0 | | 0.6 | | | | | | | | | | 0.4 |
| 22:1 | | | | | | | | | | | | |
| 24:0 | | 0.8 | 0.4 | | | | | | | | | 0.3 |
| 24:1 | | 0.7 | 0.5 | | | | | | | | | 0.3 |
| 26:0 | | | | | | | | | | | | |
| 26:1 | | | | | | | | | | | | |

n.d. High

L-mito
L-TE
C-mito
C-TE

**Supplemental Figure S3.** Fatty acid profiles of the glycerolipids from L-mito, L-TE, C-mito and C-TE. Heat maps illustrate the difference of the fatty acid distribution based on LC-MS/MS analyses. The heat map colors represent mean intensity values according to the color map on the low right-hand side. Each column represents one lipid class wherein the acyl moieties of all species are listed and summarized to 100 %. Data represent mean values in mol % from three independent experiments which were also used for proteomics (Supplemental Tab. S1).

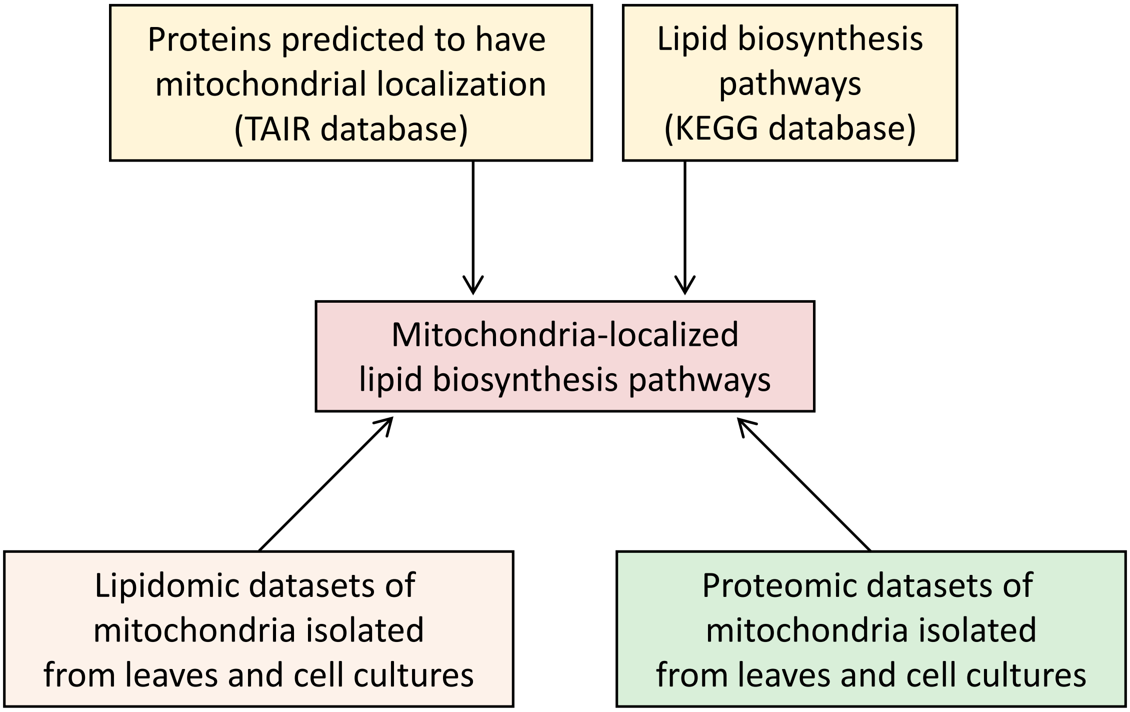


**Figure S4. Workflow for the construction of the biosynthesis pathways.**
Multiple databases were combined to build the lipid biosynthesis pathways in mitochondria, The Arabidopsis Information Resource (TAIR; www.arabidopsis.org), the Subcellular localization database for Arabidopsis proteins (SUBAcon; www.suba.live), Kyoto Encyclopedia of Genes and Genomes (KEGG; www.genome.jp/kegg) and the proteomic datasets of isolated mitochondria in this study. Information of protein localizations and backbones of the biosynthesis pathways were obtained from TAIR, SUBAcon and KEGG, respectively.

**a** **Fatty acid biosynthesis**

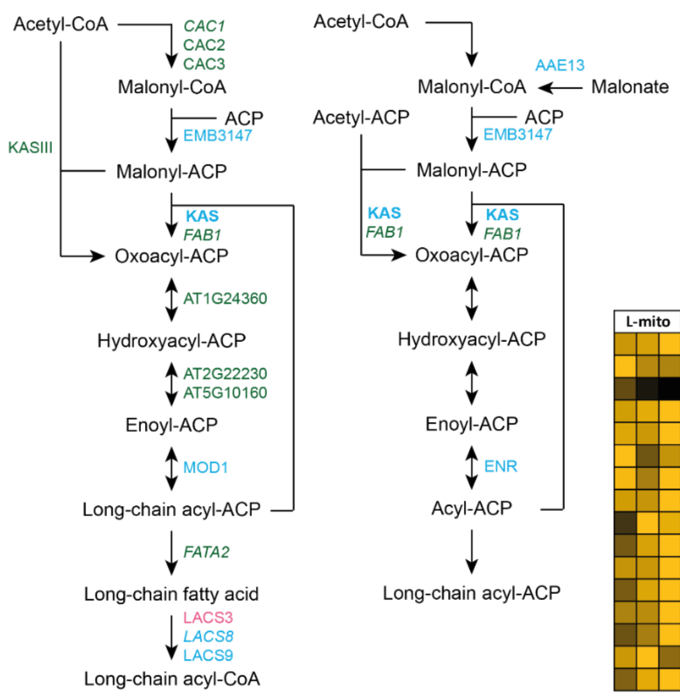


| L-mito | C-mito | Locus | Name | Function |
|---|---|---|---|---|
| | | AT1G77590 | LACS9 | Long chain acyl-CoA synthetase 9 |
| | | AT2G04350 | *LACS8* | Long-chain acyl-CoA synthetase 8 |
| | | AT2G04540 | **KAS** | 3-oxoacyl-ACP synthase |
| | | AT2G05990 | MOD1 | Enoyl-acp reductase 1 |
| | | AT2G30200 | EMB3147 | Malonyl CoA-ACP malonyltransferase |
| | | AT3G16170 | AAE13 | Acyl activating enzyme 13 |
| | | AT3G45770 | ENR | Polyketide synthase, enoylreductase |
| | | AT1G24360 | AT1G24360 | NAD(P)-binding protein |
| | | AT1G62640 | KASIII | 3-ketoacyl-ACP synthase III |
| | | AT1G74960 | *FAB1* | 3-oxoacyl-ACP synthase II |
| | | AT2G22230 | AT2G22230 | 3-hydroxyacyl-ACP dehydratase |
| | | AT2G38040 | CAC3 | Acetyl CoA carboxylase carboxyltransferase |
| | | AT4G13050 | *FATA2* | Acyl-ACP thioesterase |
| | | AT5G10160 | AT5G10160 | 3-hydroxyacyl-ACP dehydratase |
| | | AT5G16390 | *CAC1* | Chloroplastic acetyl-CoA carboxylase 1 |
| | | AT5G35360 | CAC2 | Acetyl CoA carboxylase biotin carboxylase |

**b** **Unsaturated fatty acid biosynthesis**

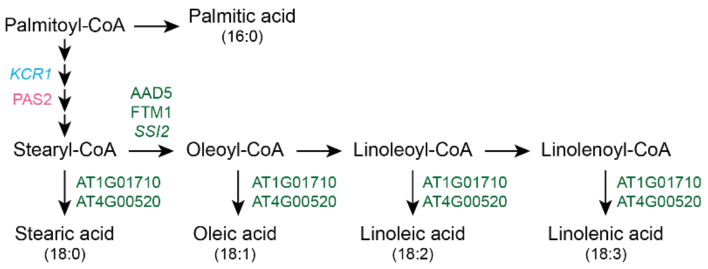


| L-mito | C-mito | Locus | Name | Function |
|---|---|---|---|---|
| | | AT1G67730 | *KCR1* | Beta-ketoacyl reductase 1 |
| | | AT1G01710 | AT1G01710 | Acyl-CoA thioesterase II |
| | | AT1G43800 | FTM1 | Stearoyl-acyl-carrier-protein desaturase |
| | | AT2G43710 | *SSI2* | Stearoyl-acyl-carrier-protein desaturase |
| | | AT3G02630 | AAD5 | Stearoyl-acyl-carrier-protein desaturase |
| | | AT4G00520 | AT4G00520 | Acyl-CoA thioesterase |

**c** **Lipoic acid metabolism**

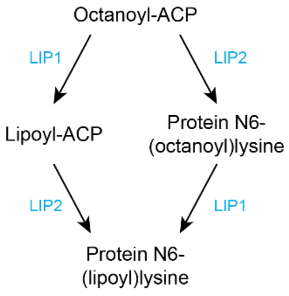


| L-mito | C-mito | Locus | Name | Function |
|---|---|---|---|---|
| | | AT1G04640 | LIP2 | Octanoyltransferase |
| | | AT2G20860 | LIP1 | Lipoyl synthase |

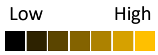

**d** **Ubiquinone and other terpenoid-quinone biosynthesis**

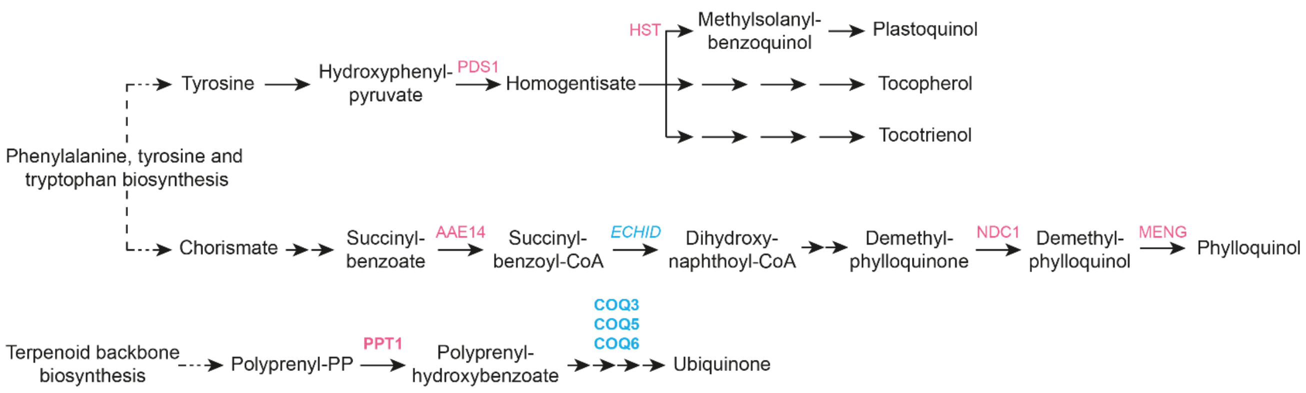


| L-mito | C-mito | Locus | Name | Function |
|---|---|---|---|---|
| | | AT1G60550 | *ECHID* | Enoyl-CoA hydratase/isomerase D |
| | | AT2G30920 | **COQ3** | Coenzyme Q 3 |
| | | AT3G24200 | **COQ6** | Ubiquinone biosynthesis monooxygenase |
| | | AT5G57300 | **COQ5** | SAM-dependent methyltransferase |

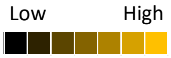


**Supplemental Fig. S5.** Additional lipid biosynthesis pathways which do not contain analyzed lipid classes in this study. Pathways of (a) fatty acid biosynthesis, (b) unsaturated fatty acid biosynthesis, (c) lipoic acid metabolism and (d) ubiquinone and other terpenoid-quinone biosynthesis illustrate the general lipid biosynthesis in Arabidopsis. Heat maps visualize the protein abundance based on shotgun proteomic analyses of three independent experiments of purified mitochondria from leaves and cell culture. The heat map colors represent mean intensity values according to the color map on the low right-hand side. Blue: proteins identified in the proteomic analysis of this study and also predicted to localize in mitochondria; green: proteins identified in the proteomic analysis of this study but predicted to localize in other organelles; magenta: proteins absent in the proteomic analysis of this study but predicted to localize in mitochondria; bold font: exclusively localized in mitochondria; italic font: only identified in one of the mitochondrial populations. Further full names and functions are itemized in Supplemental Tab. S3. In the heat maps, proteins with high and low abundance are depicted in yellow and black, respectively. Predicted protein localization was based on The Arabidopsis Information Resource (TAIR; www.arabidopsis.org) and the Subcellular localization database for Arabidopsis proteins (SUBAcon; www.suba.live).
